# Three consecutive glycolysis enzymes are involved in autophagic flux regulation through monitoring nutrient availability

**DOI:** 10.1101/2022.07.12.499818

**Authors:** Du-Hwa Lee, Ilyeong Choi, Seung Jun Park, Sumin Kim, Min-Soo Choi, Ho-Seok Lee, Hyun-Sook Pai

## Abstract

Autophagy serves as an important recycling route for growth and survival of eukaryotic organisms in nutrient-deficient conditions. When confronted with starvation, metabolic flux is coordinated by individual metabolic enzymes. Given that the metabolic diversity of carbon in eukaryotes is related to their lifestyle, autophagy may be modulated by metabolic enzymes by monitoring carbon flux. Here, we attempted to identify carbon metabolic genes that modulate autophagy using VIGS screening of 45 glycolysis- and the Calvin-Benson cycle-related genes. We report here that three consecutive triose-phosphate-processing enzymes involved in the cytosolic glycolysis, TPI (triose-phosphate-isomerase), GAPC (glyceraldehyde-3-phosphate dehydrogenase), and PGK (phosphoglycerate kinase), designated TGP, negatively regulate autophagy. Depletion of TGP enzymes result in spontaneous autophagy induction and increases ATG1 kinase activity. TGP enzymes interact with ATG101, a regulatory component of the ATG1 kinase complex. Spontaneous autophagy induction and abnormal growth under insufficient sugar in the TGP mutants is suppressed by crossing with the *atg101* mutant. Considering that triose-phosphates are photosynthates transported to the cytosol from active chloroplasts, the TGP enzymes may be strategically positioned to monitor the flow of photosynthetic sugars and modulate autophagy accordingly. Collectively, these results suggest that TGP enzymes negatively control autophagy acting upstream of the ATG1 complex, which is critical for seedling development.

## Introduction

Macroautophagy (hereafter autophagy) is a conserved catabolic process in eukaryotic cells for self-recycling through the degradation of cellular components. Autophagy is activated under starvation conditions. Eukaryotic cells employ AUTOPHAGY-RELATED (ATG) proteins for autophagic processes, from phagophore formation to fusion between autophagosomes and the lysosome or vacuole. Processing and lipidation [phosphatidylethanolamine (PE) conjugation] of ATG8 is the most fundamental step in autophagy (Thompson et al., 2005; Chung et al., 2010). Autophagic flux, which indicates cellular autophagic activity, can be assessed by the lipidation state of ATG8 (Mizushima and Yoshimori, 2007; Loos et al., 2014). In the past decade, numerous studies in mammals, using sophisticated methods, have shown that autophagic flux is tightly regulated by the mammalian Target Of Rapamycin (mTOR) and AMP-activated protein kinase pathways (Russell et al., 2014). Recent studies have suggested that the ATG1 kinase complex serves as a converging point of multiple upstream signaling pathways for the regulation of autophagic flux in *Arabidopsis*, as observed in mammals and yeast (Mizushima et al., 2010; Chen et al., 2017; Huang et al., 2019a). The ATG1 kinase complex consists of ATG1 kinase, scaffold protein ATG11, regulatory subunit ATG13, and ATG101 (Suttangkakul et al., 2011; Zhuang et al., 2018). Except that in ATG101, loss-of-function mutations in the components of the ATG1 kinase complex have been studied, and the results suggested that ATG1 kinase activity is required for autophagic body deposition under starvation conditions (Suttangkakul et al., 2011; Li et al., 2014; Huang et al., 2019a). Upon nutrient starvation, the phosphorylation status of ATG1 and ATG13 can be altered by the activity of evolutionarily conserved central metabolic regulatory kinases, Target of Rapamycin (TOR) and SNF-related kinase1 (SnRK1). ATG1 kinase controls the initiation and growth of autophagosomes, and phosphorylation/dephosphorylation of the ATG1 complex controls ATG1 activity (Suttangkakul et al., 2011; Chen et al., 2017). Collectively, these findings suggest that the upstream signaling pathways of plant autophagy and the possible regulatory mechanisms of the ATG1 complex depend on nutrient availability. However, the details are still unclear about how autophagic flux is regulated by nutrient availability in plants.

Glycolysis is an ancient metabolic pathway present in all prokaryotes and eukaryotes. Glycolysis produces pyruvate for the tricarboxylic acid cycle, ATP, reduced electron carriers, and intermediates as precursors for diverse metabolic pathways (Plaxton, 1996). As plant glycolysis occurs separately in the cytosol and plastids, more isoforms of glycolytic enzymes are present in plants than in other eukaryotes (Plaxton, 1996; Rosa-Tellez et al., 2018). Several isoforms of plastidial glycolytic enzymes, such as triose phosphate isomerase (TPI), glyceraldehyde-3-phosphate dehydrogenase (GAPDH), and phosphoglycerate kinase (PGK), are also utilized in the Calvin-Benson cycle. Both cytosolic and plastidial isoforms catalyze the same substrate in the cytosol and plastid, respectively. TPI catalyze the reversible interconversion of dihydroxyacetone phosphate (DHAP) and glyceraldehyde-3-phosphate (GAP), thereby enabling DHAP to enter glycolysis (Chen and Thelen, 2010). GAPDH catalyzes the reversible phosphorylation of GAP to 1,3-bisphosphoglycerate (1,3-BPG) in the presence of inorganic phosphate (P_i_) and nicotinamide adenine dinucleotide (NAD^+^) (Reis et al., 2013). PGK catalyzes reversible phosphoryl transfer from 1,3-BPG to ADP to produce 3-phosphoglycerate (3-PG) and ATP, the first step for ATP generation during glycolysis (Rosa-Tellez et al., 2018). The triose phosphates produced by photosynthesis are translocated to the cytosol to enter the glycolysis and sucrose biosynthesis pathways for energy production and transportation to other body parts, respectively. Thus, these series of triose-processing enzymes play an important role in carbon metabolism by connecting photosynthesis with cytosolic glycolysis and sucrose biosynthesis.

Recent studies have suggested that several glycolytic enzymes have noncanonical functions in addition to metabolic functions (Seki and Gaultier, 2017; Lu and Hunter, 2018). For example, mammalian aldolase (ALDO) regulates AMP-activated protein kinase (AMPK) activity to sense glucose availability (Zhang et al., 2017). PGK1 acts as a protein kinase for phosphorylation of Beclin1 to regulate autophagy (Qian et al., 2017). There are also examples of “moonlighting” behaviors of several glycolytic enzymes in plants. Hexokinase 1 (HXK1), an isoform of the first enzyme in the glycolysis pathway, acts as a glucose sensor and modulates the transcription of sucrose-responsive genes (Cho et al., 2007). Moreover, cytosolic GAPDH1 and 2 (GAPCs) inhibit the ROS-induced autophagy in *Nicotiana benthamiana* and *Arabidopsis* (Han et al., 2015; Henry et al., 2015). These additional functions of the enzymes, particularly the role of GAPCs in autophagy, suggest that specific glycolytic enzymes may play a role in the regulation of autophagy.

In this study, we discovered the novel functions of three specific glycolytic enzymes in the control of autophagy. The three glycolytic enzymes are cytosolic triose phosphate isomerase (CyTPI), cytosolic phosphorylating glyceraldehyde-3-phosphate dehydrogenase 1 and 2 (GAPCs), and phosphoglycerate kinase 3 (PGK3), designated TGP enzymes. The TGP enzymes catalyze the three consecutive steps in the cytosolic glycolysis pathway, which is closely connected to photosynthesis through the import of DHAP, GAP, and 3-PG from active chloroplasts (Lee et al., 2017b). We performed virus-induced gene silencing (VIGS) screening of carbon metabolic genes.

The subsequent analyses suggested that the TGP enzymes negatively regulate autophagy through modulation of ATG1 kinase activity. Considering that plants mainly produce organic carbon compounds via photosynthesis, the noncanonical functions of these glycolytic enzymes may provide a novel strategy to coordinate autophagic activity with carbon availability.

## Results

### VIGS screening to identify glycolytic enzymes that modulate autophagy

We performed VIGS screening using an *Arabidopsis* autophagy reporter line (35Sp:GFP-ATG8a) to identify negative autophagic regulators of glycolysis enzymes by monitoring autophagic flux. Since the TOR kinase complex, which consists of TOR, LST8, and RAPTOR, is a well-characterized negative regulator of autophagy (Liu and Bassham, 2010; Pu et al., 2017), we performed VIGS with *TOR*, *LST8*, and *RAPTOR* genes as proof of concept. The cDNA fragments of these genes were inserted into the pTRV2 vector, and the constructs were agroinfiltrated into the leaves of the reporter line to induce tobacco rattle virus (TRV)-mediated gene silencing. Silencing of each TOR complex gene resulted in growth retardation and premature senescence, most by *TOR* silencing (Supplemental Figure 1A). Silencing of the TOR complex genes resulted in the spontaneous formation of GFP-ATG8a puncta, approximately five times higher than that in the TRV2-myc control, suggesting that VIGS is a reliable tool to identify negative autophagic regulators (Supplemental Figure 1B, C).

We launched VIGS screening of 45 genes encoding glycolytic enzymes in *Arabidopsis* (Supplemental Figure 1D; *written in red*) and monitored GFP-ATG8a puncta levels after the silencing of each gene, as monitored in the TOR kinase complex silencing. We observed leaf epidermal cells of the VIGS plants using fluorescence microscopy and ranked them by GFP-ATG8a puncta numbers (Figure 1A). After VIGS of 45 genes, only four genes encoding TGP enzymes in the cytosolic glycolysis pathway showed statistically significant GFP-ATG8a puncta formation (Figure 1A, B). TGP enzymes include cytosolic triose phosphate isomerase (CyTPI), cytosolic phosphorylating glyceraldehyde-3-phosphate dehydrogenase 1 and 2 (GAPC1 and 2), and phosphoglycerate kinase 3 (PGK3). We performed VIGS on selected genes for confirmation of the screening results. Silencing of *CyTPI* and *PGK3* and co-silencing of *GAPC1* and *GAPC2* (*GAPCs*) resulted in retarded plant growth (Supplemental Figure 1E). The silencing efficiency of each VIGS sample was determined using quantitative reverse transcription-PCR (qRT-PCR) (Supplemental Figure 1G). The three enzymes mentioned above catalyze the sequential processing of triose phosphates, which are produced by the cytosolic glycolysis pathway, as well as by the Calvin-Benson cycle in chloroplasts and transported to the cytosolic pathway (Figure 1B). Based on the Z-stack projection of the confocal microscopy images (Figure 1C), the number of GFP-ATG8a puncta was quantified (Figure 1D). Compared with that in the control TRV2-myc, depletion of the cytosolic TGP enzymes (CyTPI, GAPCs, and PGK3) by VIGS caused statistically significant puncta formation, whereas VIGS of the plastidial TGP enzymes (PdTPI, PGK1, and PGK2) or cytosolic co-factor independent phosphoglycerate mutases (iPGAMs) had little effect (Figure 1C, D).

**Figure 1.**
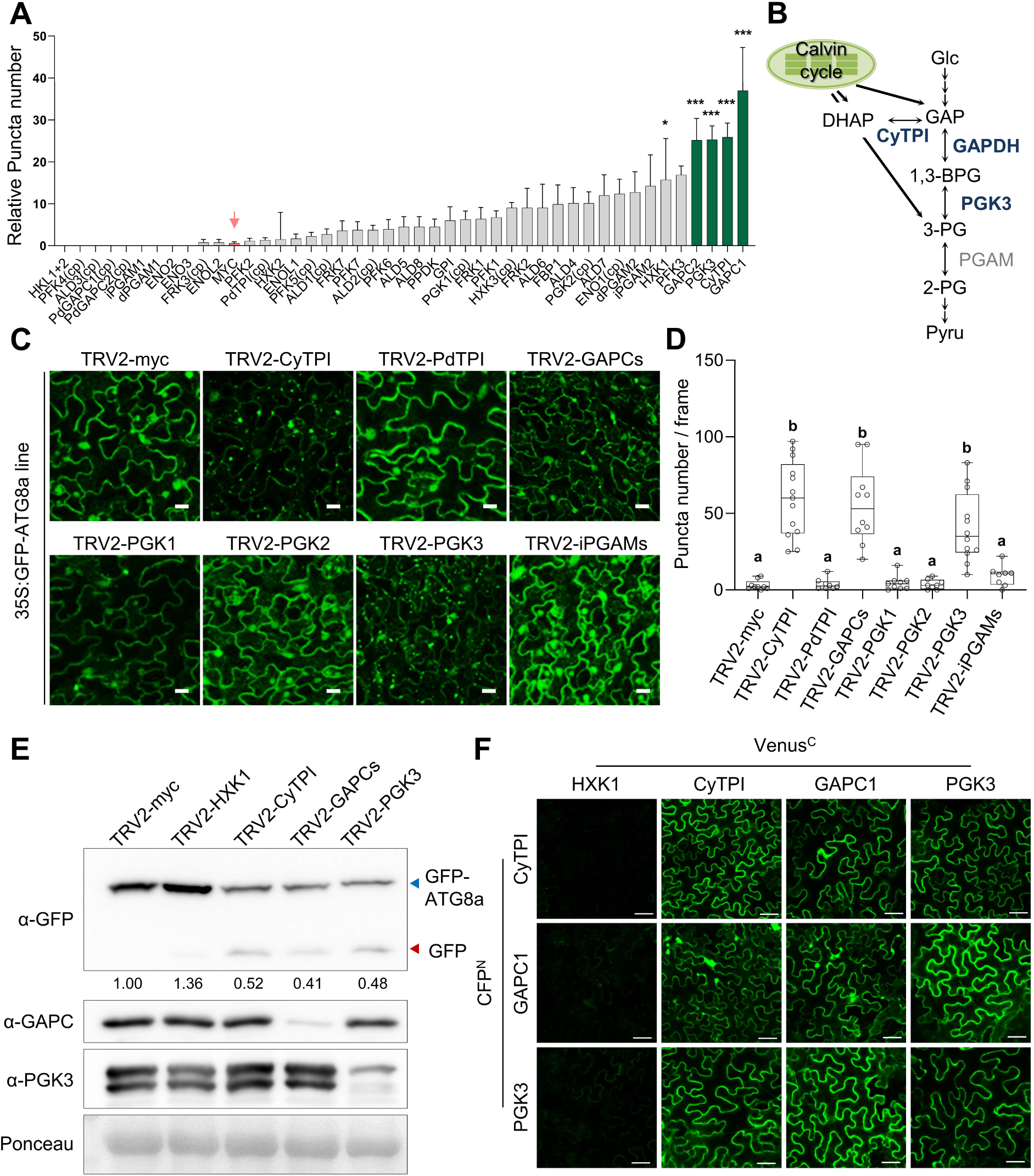
Silencing of the cytosolic TGP enzyme genes causes increased autophagic flux. (**A**) Measurement of relative puncta numbers per frame compared to TRV2-myc in fluorescent microscopic images. Enzymes are ordered from left to right by increasing relative puncta numbers. Statistical significance is assessed using linear regression analysis compared to TRV2-myc (coefficient p-value; *, p < 0.05; ***, p < 0.001). The green bars represent the genes showing statistically significant differences (p < 0.001). The pink arrow indicates TRV2-myc, the negative control of the VIGS screening. The chloroplast enzymes (including the putative ones) were marked by “(cp)” after enzyme names. (**B**) A schematic diagram of a part of the cytosolic glycolysis pathway that shows three enzymes CyTPI, GAPDH, and PGK3 (marked by *boldface*) and metabolites. Transport of triose phosphates GAP, DHAP, and 3-PG from chloroplasts to the cytosol is indicated by *thick lines*. The full glycolysis pathway in the cytosol and the Calvin-Benson cycle in chloroplasts are shown in Supplemental Figure 1D. (**C**) Confocal microscopy of GFP-ATG8a fluorescence (Z-stack image projection) in leaf epidermal cells after VIGS of the glycolytic enzyme genes at 14 days after infiltration (DAI). TRV2-myc was used as a control for VIGS. Silencing of *CyTPI, GAPCs*, and *PGK3* led to spontaneous formation of GFP-ATG8a puncta. VIGS using TRV2:GAPCs and TRV2:iPGAMs constructs caused co-silencing of *GAPC1* and *GAPC2*, and *iPGAM1* and *iPGAM2*, respectively. Scale bar = 10 μm. (**D**) Puncta numbers per frame in the Z-stack of confocal microscopy images. Dots represent puncta numbers of individual observations. The box plot represents the 1^st^ and 3^rd^ quartiles, and whiskers include the min and max of the data points. Letters above the boxes (*a* and *b*) indicate the results of one-way ANOVA followed by Tukey’s multiple comparison test (p < 0.05). (**E**) Detection of free-GFP release by autophagic breakdown of the GFP-ATG8a reporter. Total leaf extracts from VIGS plants were subjected to immunoblotting with anti-GFP, anti-GAPC, and anti-PGK3 antibodies. The GFP-ATG8a fusion protein and free-GFP are indicated by *blue* and *red arrowheads*, respectively. The anti-PGK3 antibody generated two protein bands, and the lower bands (*blue arrow*) appeared to represent PGK3 enzyme. Ponceau S-stained rubisco large subunit (rbcL) was used as the loading control. Relative band intensities of GFP-ATG8a are indicated below the anti-GFP image. (**F**) BiFC analyses for interactions between glycolytic enzymes. The CFP^N^-and Venus^C^-fusion constructs were co-expressed in *N. benthamiana* leaves via agroinfiltration. Leaf epidermal cells were observed by confocal microscopy. Scale bar = 50 μm.

The autophagic flux is frequently measured through GFP-ATG8a cleavage, which results in the production of free GFP (Li and Vierstra, 2012). Based on immunoblotting with an anti-GFP antibody, free GFP released from GFP-ATG8a was visible in VIGS samples of the TGP enzymes but not in TRV2-myc control or TRV2-Hexokinase 1 (HXK1), suggesting that a deficiency of cytosolic TGP enzymes causes an increase in autophagic flux (Figure 1E). Immunoblotting with anti-GAPC and anti-PGK3 antibodies revealed equal protein loading and reduced levels of GAPCs and PGK3 in the corresponding VIGS plants. Collectively, these data suggest that serial TGP enzymes in the cytosol may act as negative regulators of autophagy in plants.

### Cytosolic TGP enzymes interact with each other

Consecutive enzymes in a metabolic pathway frequently form enzyme subcomplexes for substrate channeling and more efficient regulation. Recent evidence suggests that the glycolytic pathway in mammalian muscle cells consists of several subcomplexes (Menard et al., 2014). To determine whether cytosolic TGP enzymes interact with each other, we performed yeast two-hybrid (Y2H) assays (Supplemental Figure 1F). GAL4 activating domain (AD)-fused CyTPI and GAL4 promoter binding domain (BD)-fused HXK1, CyTPI, GAPC1, and PGK3 were co-expressed in yeast. Based on yeast growth on selective media, CyTPI interacted with GAPC1 and PGK3, particularly strongly with CyTPI itself (Supplemental Figure 1F). However, the expression of CyTPI with the BD control or HXK1 did not support positive yeast growth, suggesting no protein interaction. To confirm the interactions between TGP enzymes *in planta*, we performed bimolecular fluorescence complementation (BiFC). CFP^N^- or Venus^C^-conjugated CyTPI, GAPC1, PGK3, and control HXK1 were expressed in combination in *N. benthamiana* leaves via agroinfiltration (Figure 1F). Every combination of CyTPI, GAPC1, and PGK3 produced green fluorescence in the cytosol, but the combination with HXK1 did not produce any fluorescent signal. Collectively, these results suggest interactions between the cytosolic TGP enzymes.

### Cytosolic TGP enzyme mutants show abnormal seedling growth depending on sucrose concentration

To confirm the results of VIGS, we obtained T-DNA insertion lines for *HXK1*, *CyTPI*, *GAPC1*, *GAPC2,* and *PGK3* (Supplemental Figure 2A). We genotyped these T-DNA insertion lines (Supplemental Figure 2B) and detected very low or undetectable levels of transcripts using qRT-PCR (Supplemental Figure 2C). Before analyzing the autophagic flux in these mutants, we observed the overall seedling growth in MS medium containing various sugar concentrations. Interestingly, the cytosolic TGP enzyme mutants *cytpi, gapc1/2* (*gapc1-1/gapc2-1* double mutant)*, pgk3-1*, and *pgk3-2* all showed similar defects in leaf development near the shoot apical meristem (SAM) under 15 mM sucrose concentration (Figure 2A; Supplemental Figure 2D). Compared to the wild-type (WT) and *hxk1-3*, these mutants produced leaves with abnormal thickness and shape, which suggested defective cell division and expansion (Figure 2A). The frequency of developmental abnormalities in the cytosolic TGP enzyme mutants was significantly higher than that in the WT and *hxk1-3* mutants; the *cytpi* and *pgk3-2* mutants exhibited particularly high levels of abnormalities (Figure 2B). However, the frequency of abnormal development in the TGP mutants significantly and progressively decreased under 30 and 60 mM sucrose (Figure 2B; Supplemental Figure 2D). These findings suggest that the deficiency of cytosolic TGP enzymes causes seedling growth defects depending on sugar concentration.

**Figure 2.**
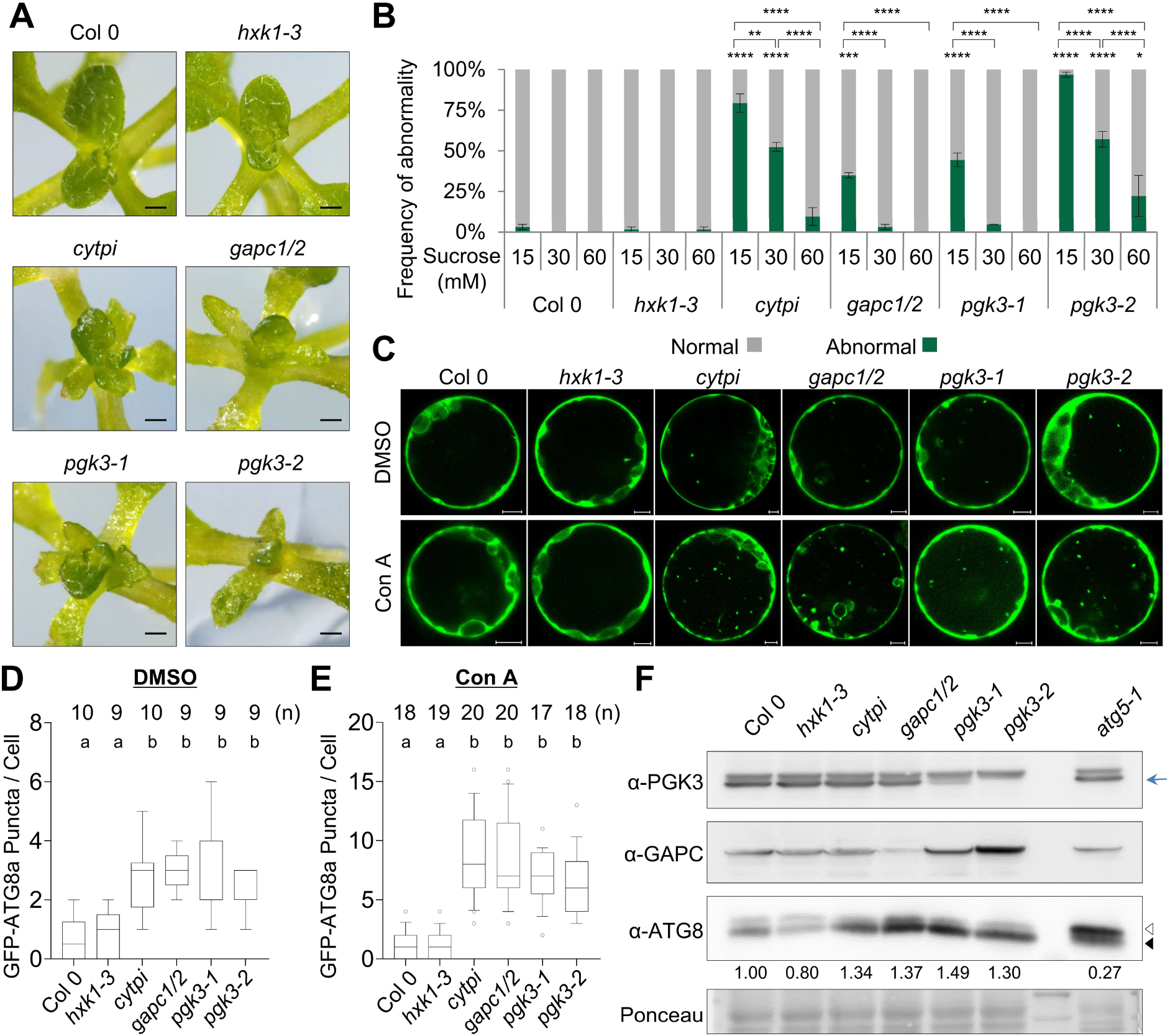
The cytosolic TGP enzyme mutants show higher autophagic flux, accompanied by abnormal shoot growth in a sugar-dependent manner. (**A**) Representative close up views of the shoot apical region in the glycolysis mutants grown in 15 mM sucrose. The whole pictures of the seedlings are shown in Supplemental Figure 2D. Emerging abnormal leaves were visible in *cytpi, gapc1/2, pgk3-1*, and *pgk3-2* mutants. Scale bars = 0.25 mm. (**B**) Frequency of abnormal shoot growth in the mutants. The bar graphs show percentages of defective seedlings out of total 20 seedlings in different sucrose concentrations (15, 30, and 60 mM). Error bars represent SE from three biological replications. Statistical significance was assessed using one-way ANOVA followed by Tukey’s multiple comparison test (*, p < 0.05; **, p < 0.01; ***, p < 0.001; ****, p < 0.0001). The statistical significances compared to WT (Col-0) under the same sucrose concentration are marked at the top of the bars, while other comparisons are indicated by respective *brackets*. (**C**) Confocal microscopy of GFP-ATG8a fluorescence (Z-stack image projection) in *Arabidopsis* mesophyll protoplasts of the glycolysis mutants. Leaf protoplasts were transfected with the GFP-ATG8a construct, followed by treatment with DMSO (control) or 1 μM concanamycin A (Con A). Scale bars = 5 μm. (**D and E**) The GFP-ATG8a puncta in the protoplasts were counted in the Z-stacked confocal microscopy images after DMSO (**D**) or Con A (**E**) treatment. Puncta numbers per frame are shown in the Z-stack of confocal microscopy images. The box plot represents the 1^st^ and 3^rd^ quartiles, and whiskers include the 10^th^-90^th^ percentile of the data points. The number of biologically independent observations (n) is indicated at the top of the graph. Letters above the boxes (*a-c*) indicate the results of one-way ANOVA followed by Tukey’s multiple comparison test (p < 0.05). (**F**) Immunoblotting for monitoring ATG8-PE/ATG8 patterns in the glycolysis mutants. Twelve-day-old seedlings were incubated in the light in liquid medium containing 30 mM glucose. Protein extracts were subjected to immunoblotting with anti-PGK3, anti-GAPC, and anti-ATG8 antibodies. The atg5-1 autophagy mutant was used as the positive control. The ATG8 and ATG8-PE are marked by *open* and *closed arrowheads*, respectively. The PGK3 protein band is marked by the *blue arrow*. Relative band intensities of ATG8-PE/ATG8 are indicated below the anti-ATG8 image.

### Cytosolic TGP enzyme mutants have higher autophagic flux than wild-type

To examine autophagic flux in the cytosolic TGP enzyme mutants, we observed autophagosomes by following the fluorescence of GFP-ATG8a in WT and the glycolytic mutants using confocal microscopy (Figure 2C). Mesophyll protoplasts were prepared from the leaves of four-week-old WT and mutant plants grown in soil and transfected with the GFP-ATG8a construct. The protoplasts were then treated with the dimethyl sulfoxide (DMSO) control or Concanamycin A (Con A) for 16 h. Con A blocks the vacuolar degradation of autophagosomes by specifically inhibiting vacuolar ATPase (Dettmer et al., 2006). GFP-ATG8a puncta in protoplasts from various lines were counted using Z-stack image projection with confocal microscopy (Figure 2D, E). The average number of fluorescent puncta in the protoplast was higher in the cytosolic TGP enzyme mutants than in the WT and *hxk1-3* mutants, and the effect was more visible upon Con A treatment (Figure 2C-E). Increased puncta numbers following Con A treatment suggested that the formation of excessive puncta in the TGP enzyme mutants was not caused by vesicle trafficking defects. These observations, i.e. the increase in ATG8a-positive puncta formation in the cytosolic TGP enzyme mutants, were consistent with the results of VIGS screening (Figure 1C, D).

To confirm the increased autophagic flux in the cytosolic TGP enzyme mutant, we examined the band patterns of ATG8-PE and ATG8 in the mutants using the autophagy mutant *atg5-1* as a control. Immunoblotting using anti-PGK3 and anti-GAPC antibodies revealed reduced protein levels in their corresponding mutants, with Ponceau S-stained rbcL (Rubisco large subunit) as the loading control (Figure 2F). Immunoblotting using an anti-ATG8 antibody showed that the ATG8-PE form (*closed arrowhead*) was more abundant than the free-ATG8 form (*open arrowhead*) in the cytosolic TGP enzyme mutants compared to that in the WT and *hxk1-3* mutants not only after dark/starvation treatment for 2 days (Supplemental Figure 2E), but also under normal conditions (Figure 2F). The control *atg5-1* mutant accumulated a large amount of the free-ATG8 form, which suggested significant blockage of autophagic flux. Collectively, these results suggest that the depletion of cytosolic TGP enzymes increases the autophagic flux.

### Cytosolic TGP enzymes interact with ATG101

Upon nutrient starvation, energy signaling pathways such as the TOR and SnRK1 pathways converge on the ATG1 kinase complex to modulate autophagic flux (Chen et al., 2017; Pu et al., 2017). As autophagic flux is directly related to ATG1 kinase activity in eukaryotes, we examined the possible interactions of glycolytic enzymes, including cytosolic TGP enzymes, with the regulatory subunits of the ATG1 kinase complex (ATG101, ATG13a, and ATG13b). To determine these interactions *in planta*, we performed BiFC assays. Venus^N^-fused glycolytic enzymes (HXK1, CyTPI, PdTPI, GAPC1, GAPC2, PGK2, and PGK3) and Venus^C^-fused ATG101 were co-expressed in *N. benthamiana* leaves via agroinfiltration. Confocal microscopy detected Venus fluorescence in the cytosol of leaf epidermal cells expressing CyTPI, GAPC1, GAPC2, and PGK3, along with ATG101, suggesting interactions between ATG101 and these enzymes. Among these positive interactions, large fluorescent foci were observed with CyTPI, GAPC1, and GAPC2 but not with PGK3 (Figure 3A, *upper*). These BiFC results were confirmed by fusing the constructs with Venus^N^ or Venus^C^ in a reverse manner (Figure 3A, *lower*). Both BiFC experiments consistently showed that ATG101 interacts with GAPC1, GAPC2, CyTPI, and PGK3, but not with PdTPI and PGK2, and only weakly with HXK1. In addition, we performed BiFC with ATG13a and ATG13b, which are the regulatory components of the ATG1 kinase complex. The Venus^C^-fused GAPC1, GAPC2, CyTPI, and PGK3 yielded relatively low levels of fluorescence when co-expressed with the Venus^N^-fused ATG13a and ATG13b (Supplemental Figure 3A). However, the expression of ATG13a or ATG13b in the presence of HXK1, PdTPI, or PGK2 did not generate fluorescence. These results suggest that cytosolic TGP enzymes interact with ATG101 and ATG13 in the cytosol as well as in some foci.

**Figure 3.**
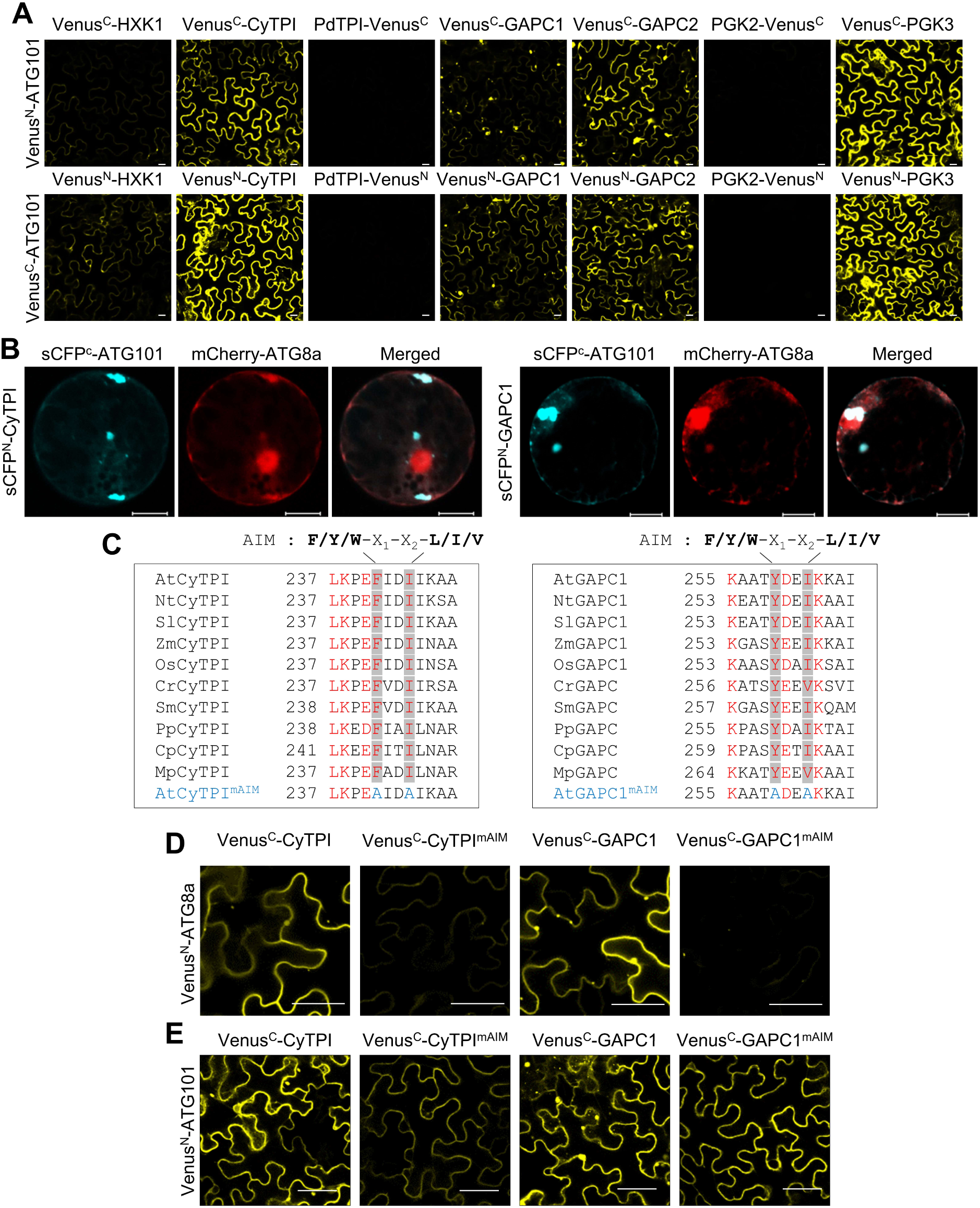
The cytosolic TGP enzymes interact with ATG101. (**A**) BiFC. Combinations of VENUS^N^- and VENUS^C^-fusion proteins were co-expressed in *N. benthamiana* leaves via agroinfiltration. Venus^N^ and Venus^C^ were fused to the C-terminus of PdTPI and PGK2 that are targeted to chloroplasts. Leaf epidermal cells were observed by confocal microscopy. Scale bars = 20 μm. (**B**) Subcellular co-localization of BiFC fluorescence (*blue*) and mCherry-ATG8a fluorescence (*red*) in leaf protoplasts. Combinations of CFP^N^- and CFP^C^-fusion proteins were co-expressed with mCherry-ATG8a in *N. benthamiana* leaves via agroinfiltration. Scale bar = 20 μm. (**C**) Conserved AIM sequences in CyTPI and GAPC1 from various plant species. Nt, *Nicotiana tabacum*; Sl, *Solanum lycopersicum*; Zm, *Zea mays*; Os, *Oryza sativa*; Cr, *Ceratopteris richardii* (fern); Sm, *Selaginella moellendorffii* (lycophytes), Pp, *Physcomitrium patens* (bryophytes); Cp, *Ceratodon purpureus* (bryophytes); and Mp, *Marchantia polymorpha* (bryophytes). The consensus sequence of the AIM is shown above the boxes. The first amino acid residue number of the AIM sequence is shown in each protein. Both AtCyTPI^mAIM^ and AtGAPC1^mAIM^ have two amino acid changes to alanine in the conserved AIM sequences as indicated. (**D and E**) BiFC analyses to detect interactions of the glycolytic enzymes and their mutants (mAIM) with ATG8a (**D**) and ATG101 (**E**). VENUS^N^- and VENUS^C^-fusion proteins were co-expressed in *N. benthamiana* leaves by agroinfiltration. Leaf epidermal cells were observed by confocal microscopy. Scale bar = 20 μm.

To confirm the interactions between cytosolic TGP enzymes and ATG101, we performed Y2H assays (Supplemental Figure 3B). AD-fused glycolytic enzymes and BD-fused ATG101 were co-expressed in yeast. Positive yeast growth in selective media suggested interactions between TGP enzymes and ATG101, whereas combinations of HXK1 and ATG101 or AD control and ATG101 did not promote yeast growth, i.e., no protein interaction (Supplemental Figure 3B), consistent with the results from BiFC (Figure 3A). Finally, we performed co-immunoprecipitation experiment to detect protein interactions. HA-tagged glycolytic enzymes (HXK1, CyTPI, GAPC1, and PGK3) and Myc-tagged ATG101 were expressed together in *N. benthamiana* leaves via agroinfiltration. Myc-ATG101 was immunoprecipitated using an anti-Myc antibody-conjugated resin. Immunoblotting with an anti-HA antibody revealed that HA-CyTPI, HA-GAPC1, and HA-PGK3 co-immunoprecipitated with Myc-ATG101 but not with HA-HXK1 (Supplemental Figure 3C). Collectively, these results suggest that CyTPI, GAPCs, and PGK3 interact with ATG101 *in vivo* and that cytosolic TGP enzymes may be involved in autophagic flux regulation.

### CyTPI and GAPC1 interact with ATG101 at the foci through ATG8

Phosphatidylethanolamine (PE)-conjugated ATG8 is associated with the autophagosomal membrane (Li and Vierstra, 2012). We tested whether the large foci structures observed in the BiFC assays (Figure 3A) co-localized with ATG8. CFP^C^-ATG101 was co-expressed with CFP^N^-fused CyTPI and GAPC1 and with mCherry-ATG8a in *N. bethamiana* leaves via agroinfiltration. Protoplasts were then isolated from the infiltrated leaves for observation under a confocal microscope at different excitation wavelengths. Co-expression of CFP^C^-ATG101 with CFP^N^-CyTPI or CFP^N^-GAPC1 caused the formation of fluorescent foci-like structures that colocalized with the red fluorescence of mCherry-ATG8a, at least partially (Figure 3B).

As ATG8-interacting proteins generally contain the ATG8-Interacting motif (AIM), we analyzed the amino acid sequences of CyTPI, GAPC1, GAPC2, PGK3, and ATG101 to search for the consensus sequence of AIM (F/Y/W-X_1_-X_2_-L/I/V) (Jia et al., 2019; Marshall et al., 2019). Interestingly, the AIM consensus sequence was found in CyTPI, GAPC1, and GAPC2 but not in PGK3 or ATG101 (Supplemental Figure 3D), suggesting that ATG8 interactions with these enzymes may play a role in the formation of the large fluorescent foci. Interestingly, the AIM sequences in the CyTPI and GAPC1 proteins were conserved in higher and lower plant species, including dicots, monocots, ferns, lycophytes, and bryophytes (Figure 3C). We constructed CyTPI and GAPC1 variants with disrupted AIM sequences. CyTPI^mAIM^ contained a double mutation of F241A and I244A, whereas GAPC1^mAIM^ contained a double mutation of Y259A and I262A. The variants fused with Venus^C^ were co-expressed with Venus^N^-fused ATG8a in *N. benthamiana* leaves for BiFC. Confocal microscopy for Venus fluorescence revealed that wild-type CyTPI and GAPC1 interacted with ATG8a in the cytosol and some foci (Figure 3D). However, BiFC of CyTPI^mAIM^ and GAPC1^mAIM^ variants with ATG8a did not generate fluorescence, suggesting that AIM sequences are critical for their interactions (Figure 3D). BiFC assays also indicated that wild-type CyTPI and GAPC1 could interacted with ATG101 in the cytosol and foci, but their variants containing the mAIM sequences interacted with ATG101 only in the cytosol (Figure 3E). Collectively, these results suggest that CyTPI and GAPC1 require their AIMs to interact with ATG8a and that the AIMs are critical for interactions between CyTPI/GAPC1 and ATG101 in large foci.

### Loss-of-function mutation of *ATG101* results in plant phenotypes similar to those of other ATG1 kinase complex subunit mutations

The ULK1/2 kinase complex, the mammalian homolog of the ATG1 complex, regulates autophagic flux during nutrient starvation (Russell et al., 2013). In plants, the ATG1 complex is critical for autophagosome enclosure and/or vacuolar delivery (Suttangkakul et al., 2011). *Arabidopsis* ATG1 kinase complex consists of ATG1, ATG11, ATG13, and ATG101; the functions of these components in autophagic processes have been characterized, except for those of ATG101. The *atg11* single mutant, *atg13a/atg13b* double mutant, and *atg1abct* quadruple mutant showed typical autophagy-deficient phenotypes, such as hypersensitivity to nitrogen or carbon starvation and premature leaf senescence under short-day conditions (Suttangkakul et al., 2011; Li et al., 2014; Huang et al., 2019a). We obtained a mutant line with a T-DNA insertion in the 3^rd^ intron of *ATG101*, designated as *atg101-1* (Supplemental Figure 4A, B). qRT-PCR revealed reduced *ATG101* transcript levels in the mutant plants compared to those in the WT plants (Supplemental Figure 4C). We next generated two independent *atg101-1* complementation lines, designated *atg101-1* C-13 and *atg101-1* C-14, by transforming the *atg101-1* mutant plant with a construct containing the *YFP-ATG101* gene under the control of its own promoter (*ATG101p:YFP-ATG101*).

We performed a carbon-starvation recovery experiment to examine the phenotypes of the *atg101-1* mutant and its complementation lines (C13 and C14) compared to those of the WT (Col-0 and Col-3) and other autophagic mutants, *atg13a/atg13b*, *atg11-1*, and *atg5-1*. Twelve-day-old seedlings grown under light/glucose (LG) conditions were transferred to dark/starvation (DS) conditions. After 5 days of incubation in DS, the seedlings were transferred to LG conditions (ReLG) and incubated for 5 days to monitor the re-greening phenotype. The WT seedlings (Col-0 and Col-3) generated green leaves after 5 days of ReLG treatment, whereas *atg101-1*, *atg13a/atg13b*, *atg11-1*, and *atg5-1* mutants did not produce green leaves (Figure 4A). The *atg101-1* complementation lines (C13 and C14) showed re-greening phenotypes similar to those of the WTs, suggesting that the *atg101-1* mutant phenotype was rescued by expression of the *YFP-ATG101* gene (Figure 4A). The total chlorophyll content was remarkably different between these seedlings, which perfectly matched the visible phenotypes (Figure 4A, C). We further examined plant responses to nitrogen starvation. Ten-day-old seedlings grown under normal conditions were transferred to either nitrogen-rich (N+) or nitrogen-deficient (N −) media and incubated for 5 days (Figure 4B). All seedlings were severely affected by nitrogen deficiency, but the *atg101-1*, *atg13a/atg13b*, *atg11-1*, and *atg5-1* mutants contained much lower amounts of chlorophyll than the WT and complementation lines, suggesting more severe defects in autophagy mutants (Figure 4B, D).

**Figure 4.**
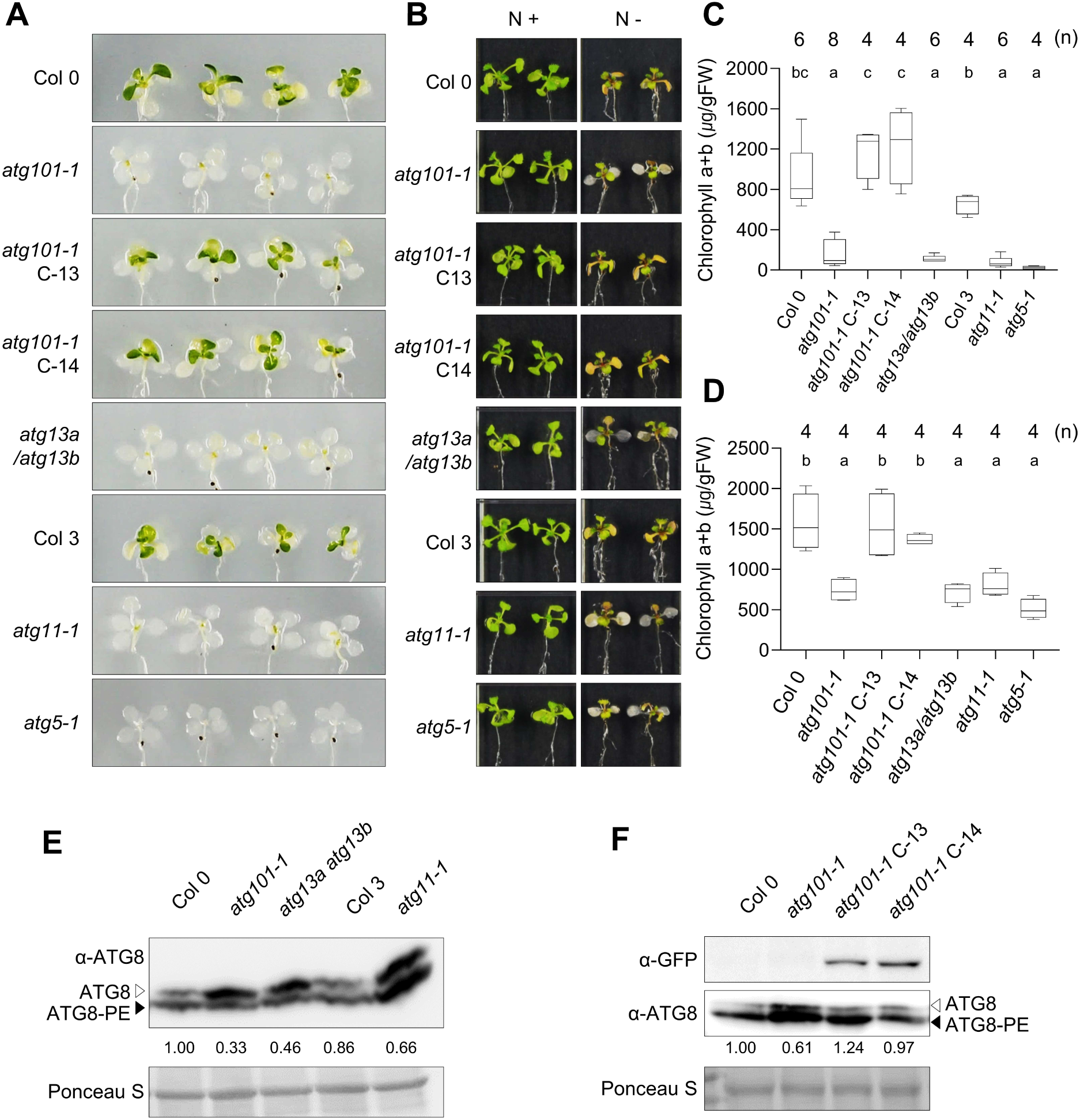
The *atg101-1* mutant shows the typical autophagy mutant phenotype and decreased autophagic flux. (**A**) Carbon starvation phenotypes of the ATG1 complex mutants and their complementation lines. Twelve-day-old seedlings were transferred to fresh liquid medium lacking glucose and incubated in the dark for 5 days. Subsequently, the seedlings were supplied with 30 mM glucose and grown in the light for 5 days. The *atg101-1* C-13 and C-14 are two complementation lines of the *atg101-1* mutant, expressing *ATG101p:YFP-ATG101*. The *atg5-1* autophagy mutant was used as the positive control. (**B**) Nitrogen starvation phenotype of the ATG1 complex mutants. Ten-day-old seedlings were transferred to media with nitrogen (N+) or without nitrogen source (N-), and incubated for 5 days in the light. (**C and D**) Total chlorophyll contents in the seedlings shown in (**A**) and (**B**). The box plots represent the 1^st^ and 3^rd^ quartiles, and whiskers include the 10^th^-90^th^ percentile of the data points. The number of biologically independent observations (n) is marked on the top of the graph. Letters above the boxes (*a-c*) indicate the results of one-way ANOVA followed by Tukey’s multiple comparison test (p < 0.05). (**E and G**) Immunoblotting for monitoring ATG8-PE:ATG8 patterns in the ATG1 complex mutants and the atg101 complementation line. Twelve-day-old seedlings were subjected into immunoblotting with anti-ATG8 antibody. The ATG8 and ATG8-PE are marked by *open* and *closed arrowheads*, respectively. Ponceau S-stained rubisco large subunit (rbcL) was used as the loading control. Relative band intensities of ATG8-PE/ATG8 are indicated below the anti-ATG8 image.

Next, we analyzed autophagic flux via immunoblotting to visualize the ATG8-PE and free-ATG8 forms. Immunoblotting using an anti-ATG8 antibody showed that the free-ATG8 form (*closed arrowhead*) was more abundant in the *atg101-1* mutant than in the WT, as similarly observed in other ATG1 complex mutants (*atg13a/atg13b* and *atg11-1*) (Figure 4E). In contrast, the abundance of the free-ATG8 form returned to the WT level in the complementation lines (*atg101-1* C13 and C14) (Figure 4F). The anti-YFP antibody detected the expression of YFP-ATG101 in the atg101 C13 and C14 lines. Next, we observed and quantified ATG8a-positive autophagosomes in *atg101-1*, *atg101-1* C13, *atg101-1* C14, and *atg13a/atg13b* lines, compared to those in the WT (Col-0) (Supplemental Figure 4D, E). Mesophyll protoplasts prepared from various lines were transfected with the GFP-ATG8a construct and treated with Con A for 16 h. GFP-ATG8a puncta in the protoplasts were counted using Z-stack image projection with confocal microscopy (Supplemental Figure 4D, E). The average number of fluorescent puncta was lower in *atg101-1* and *atg13a/atg13b* mutants than in the WT and *atg101-1* C13 and *atg101-1* C14, suggesting reduced autophagic flux in the autophagic mutants (Supplemental Figure 4E). Collectively, these results revealed the function of ATG101; its deficiency delayed the progression of autophagy, as observed with other mutations of the ATG1 complex.

### Crossing with the *atg101-1* mutant alleviates the growth defects of the cytosolic TGP enzyme mutants

Since cytosolic TGP enzymes interact with ATG101 (Figure 3), we crossed glycolytic enzyme mutants with the *atg101-1* mutant to determine a possible genetic relationship. Seedlings were observed for abnormality in growth under various sugar concentrations (Figure 5A; Supplemental Figure 5). Both WT and *atg101-1* seedlings showed normal growth in media containing 15 mM sucrose, as was also observed in the *hxk1-3* and *hxk1-3 atg101-1* double mutants (Figure 5A, 5B; Supplemental Figure 5). As shown in Figure 2, a high percentage of the TGP mutant seedlings exhibited abnormal leaves near the SAM with altered thickness and shape in 15 mM sucrose (Figure 5B). *pgk3-2* seedlings showed a particularly high percentage of developmental defects.

**Figure 5.**
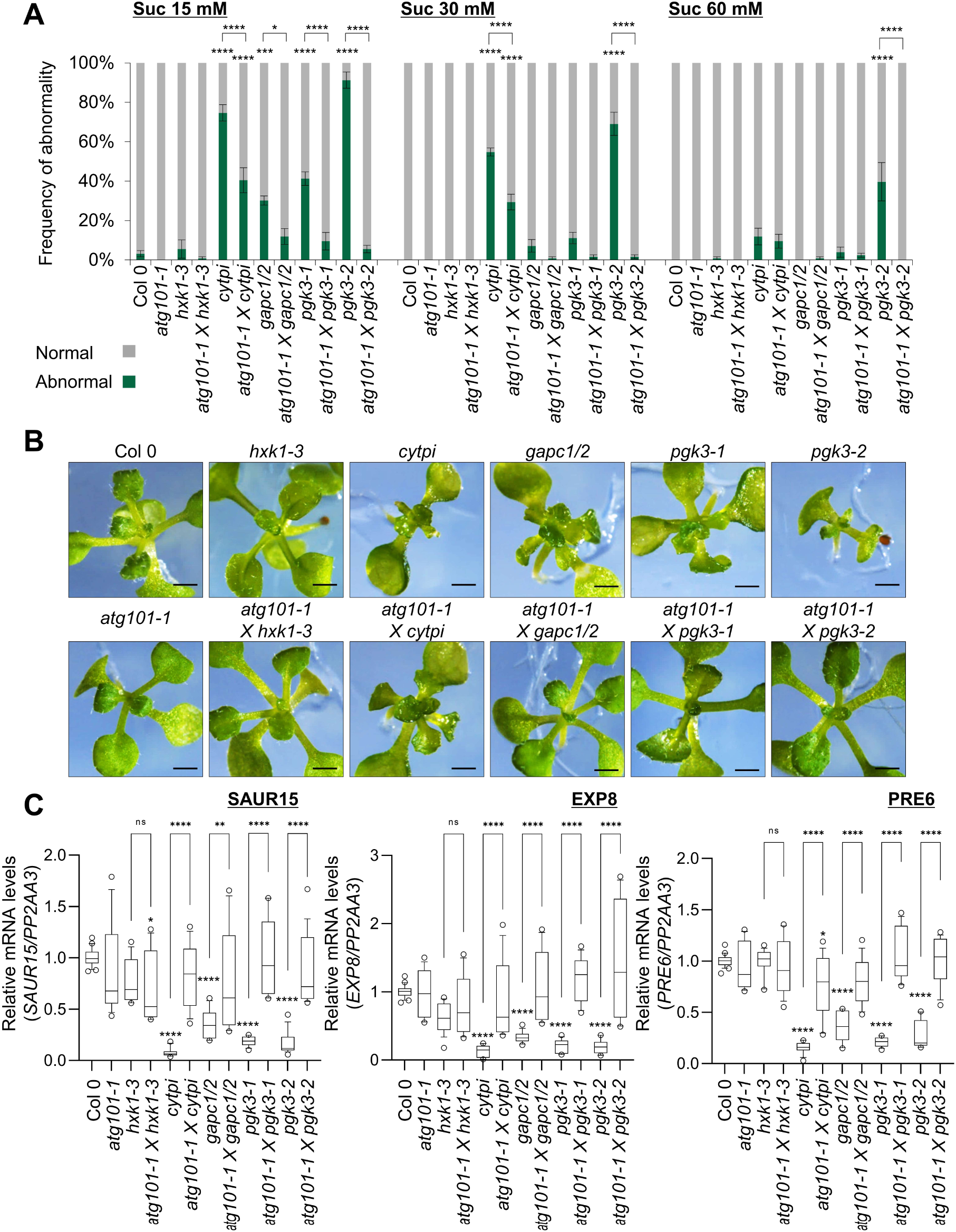
Abnormal growth phenotype of the glycolytic enzyme mutants is rescued by crossing with the *atg101-1* mutant. (**A**) Frequency of abnormality in the seedlings. The bar graphs show percentages of abnormal seedlings out of total 20 seedlings in 15 mM, 30mM, and 60 mM sucrose. Error bars represent SE from biological replications (n = 6). Statistical significance was assessed using one-way ANOVA followed by Tukey’s multiple comparison test (*, p < 0.05; ***, p < 0.001; ****, p < 0.0001). The statistical significances compared to WT (Col-0) are marked at the top of the bars, while other comparisons are indicated by the respective *brackets*. (**B**) Representative close up views of the shoot apical regions of the seedlings shown in 15 mM sucrose. Scale bar = 0.2 mm. (**C**) qRT-PCR analyses of cell expansion-related genes in the seedlings shown in 15 mM sucrose. The box plots represent the 1^st^ and 3^rd^ quartiles, and whiskers include the 10^th^-90^th^ percentile of the data points. Statistical significance was assessed from 4 biological replications using one-way ANOVA followed by Tukey’s multiple comparison test (*, p < 0.05; **, p < 0.01; ****, p < 0.0001).

However, increasing sucrose concentrations (30 and 60 mM) progressively suppressed the abnormal growth of TGP mutants, including *pgk3-2* (Figure 5A; Supplemental Figure 5). Interestingly, crossing with the *atg101-1* mutant significantly suppressed abnormal leaf growth in these mutants under all sucrose conditions (Figure 5A; Supplemental Figure 5). In addition, qRT-PCR demonstrated that the expression of *SAUR15*, *EXP8*, and *PRE6*, which are cell growth-related genes under the control of auxin and brassinosteroid, was downregulated in the cytosolic TGP enzyme mutants but significantly increased upon crossing with the *atg101-1* mutant, consistent with their improved growth (Figure 5A; Supplemental Figure 5). Combined with the increased autophagic flux in the mutants (Figure 2), these genetic interactions suggested that cytosolic TGP enzymes may play a role in plant autophagy, possibly by modulating the activity of the ATG1 complex. Interactions with ATG101 may contribute to the noncanonical function of TGP enzymes.

### Deficiency of ATG101 suppresses the elevated autophagic flux in the cytosolic TGP enzyme mutants

To determine whether altered autophagic flux contributes to the abnormal growth of cytosolic TGP enzyme mutants, we assessed autophagic flux in cytosolic TGP enzyme mutants and *atg101-1*-crossed double mutants. Twelve-day-old seedlings grown in LG were transferred to LG (as control) or DS for 2 days of incubation and then subjected to immunoblotting with anti-PGK3, anti-GAPC, and anti-ATG8 antibodies. Anti-PGK3 and anti-GAPC antibodies detected reduced protein levels of PGK3 and GAPC in *pgk3* (*pgk3-1* and *pgk3-2*) and *gapc1/2* mutants, respectively, in LG (Figure 6A) and DS (Supplemental Figure 6) conditions. All DS samples accumulated higher levels of ATG8-PE than the LG samples, which indicated that two days of dark/starvation treatment could induce autophagy (Figure 6A; Supplemental Figure 6). The ATG8-PE adduct levels were higher in the cytosolic TGP nzyme mutants *cytpi*, *pgk3-1*, *pgk3-2*, and *gapc1/2* than in the WT and *hxk1-3* mutants under LG and DS conditions, suggesting an increased autophagic flux due to the deficiency of TGP enzymes (Figure 6A; Supplemental Figure 6), consistent with previous data (Figure 2F; Supplemental Figure 2E). Introduction of the *atg101-1* mutation suppressed the increase in ATG8-PE levels in the TGP enzyme mutants, indicating that ATG101 plays a role in increasing the autophagic flux of TGP enzyme mutants. The patterns of ATG8-PE:ATG8 revealed blockage of autophagic flux in the control *atg5-1* mutant (Figure 6A).

**Figure 6.**
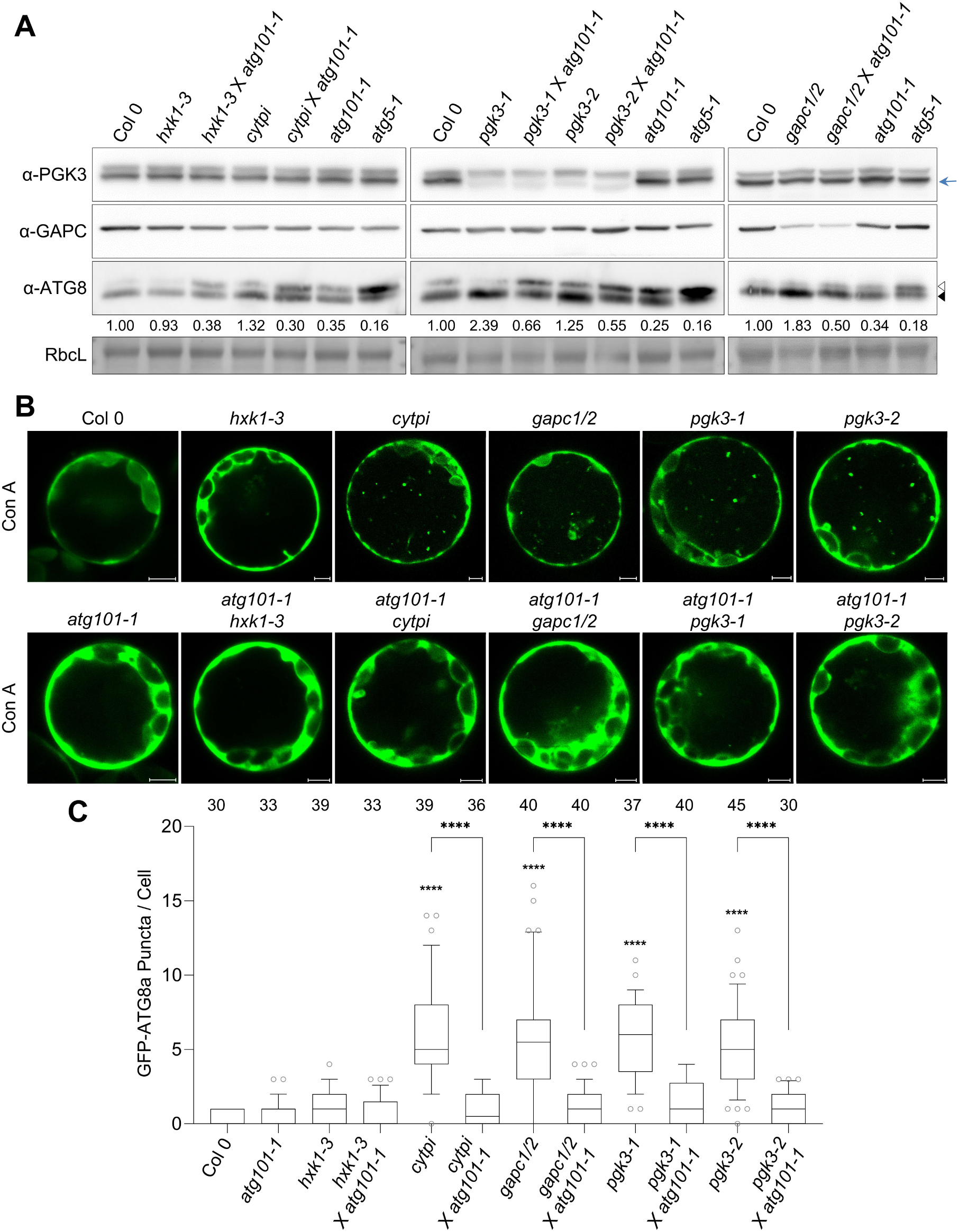
Deficiency of ATG101 suppresses increased autophagy flux in the glycolytic enzyme mutants. (**A**) Immunoblotting to examine ATG8-PE:ATG8 patterns in the single glycolysis mutants and the double mutants (*atg101-1* cross lines). Twelve-day-old seedlings were transferred to liquid medium containing 30 mM glucose and incubated in the light for 2 days. Then immunoblotting was performed with anti-PGK3, anti-GAPC1, and anti-ATG8 antibodies. Ponceau S-stained rubisco large subunit (rbcL) was used as the loading control. The ATG8 and ATG8-PE are marked by *open* and *closed arrowheads*, respectively. The PGK3 protein band is marked by the *blue arrow*. Relative band intensities of ATG8-PE/ATG8 are indicated below the anti-ATG8 image. (**B**) Observation of the GFP-ATG8a fluorescence (Z-stack image projection) in the mesophyll protoplasts of the single and double mutants. Leaf protoplasts were transfected with the GFP-ATG8a construct, followed by Con A (1 µM) treatment. Then GFP-ATG8a puncta in the protoplasts were observed under a confocal microscope. Scale bars = 5 μm. (**C**) The GFP-ATG8a puncta in the protoplasts were counted in the Z-stacked confocal microscopy images after Con A treatment. The box plot represents the 1^st^ and 3^rd^ quartiles, and whiskers include the 10^th^-90^th^ percentile of the data points. The number of biologically independent observations (n) is indicated on top of the graph. Statistical significance was eveluated using one-way ANOVA followed by Tukey’s multiple comparison test (****, p < 0.0001). The statistical significances compared to WT (Col-0) are marked at the top of the bars, while other comparisons are indicated by the respective *brackets*.

Next, we observed autophagosomes by visualizing GFP-ATG8a puncta in leaf mesophyll protoplasts of the WT, TGP enzyme mutants, and their *atg101-1* cross lines (Figure 6B). Protoplasts were transfected with the GFP-ATG8a construct, followed by 16 h of treatment with Con A. GFP-ATG8a puncta were counted using Z-stack image projection of confocal microscopy. The number of fluorescent puncta was higher in the cytosolic TGP enzyme mutants than in their *atg101-1* cross lines and WT (Figure 6B, C), which is consistent with the elevation of ATG8 lipidation in the mutants (Figure 6A; Supplemental Figure 6A). Thus, the *atg101* mutation appears to suppress the formation of ATG8-PE adducts and autophagosomal puncta in TGP enzyme mutants. Taken together, these results suggest that cytosolic TGP enzymes can modulate autophagic flux by acting upstream of the ATG1 complex, at least partly through the ATG101 interaction.

### Cytosolic TGP enzymes negatively regulate ATG1 kinase activity

Based on the genetic and biochemical relationship between cytosolic TGP enzymes and ATG101, we hypothesized that stimulation of autophagic flux in cytosolic TGP enzyme mutants may be caused by an increase in ATG1 kinase activity. To address this hypothesis, we measured the kinase activity of ATG1 *in planta*. Plant ATG6/Beclin/Vps30 (ATG6) is a subunit of the phosphatidylinositol-3-kinase (PI3K) complex that functions downstream of the ATG1 kinase complex (Liu et al., 2005; Huang et al., 2019b). Beclin, the mammalian homolog of ATG6, is a well-characterized downstream substrate of ULK1/2 (Russell et al., 2013). We performed kinase assays using immunoprecipitated YFP-ATG1a with purified recombinant MBP or MBP-ATG6 as the substrate (Supplemental Figure 7A). We first demonstrated the autophosphorylation activity of YFP-ATG1a via autoradiography of a phosphorylated protein band of the correct size (Supplemental Figure 7B). We then examined whether YFP-ATG1a possesses transphosphorylation activity towards MBP-ATG6. The kinase assay showed that MBP-ATG6, but not MBP, was phosphorylated by YFP-ATG1a kinase, suggesting that ATG6 is a substrate of ATG1a in plants, findings similar to those reported in mammals (Supplemental Figure 7C, D). Notably, when MBP-ATG6 was present, MBP-ATG6 transphosphorylation mostly occurred, whereas ATG1a autophosphorylation significantly decreased (Supplemental Figure 7D). Next, to examine whether ATG1 kinase activity is related to the autophagic flux of cytosolic TGP enzymes, we performed kinase assays using immunoprecipitated YFP-ATG1a from VIGS plants (*35Sp:YFP-ATG1a* background) of the TGP enzymes and control genes (Supplemental Figure 7E). Based on immunoblotting and quantification, the TGP VIGS samples showed increased phospho-band intensity in both ATG1a autophosphorylation and ATG6 phosphorylation compared with that in the TRV2-myc control (Figure 7A-D). Thus, the deficiency of cytosolic TGP enzymes stimulates ATG1 kinase activity, subsequently increasing autophagic flux. Taken together, these data suggest that cytosolic TGP enzymes negatively regulate autophagy by modulating ATG1 kinase activity (Figure 7E).

**Figure 7.**
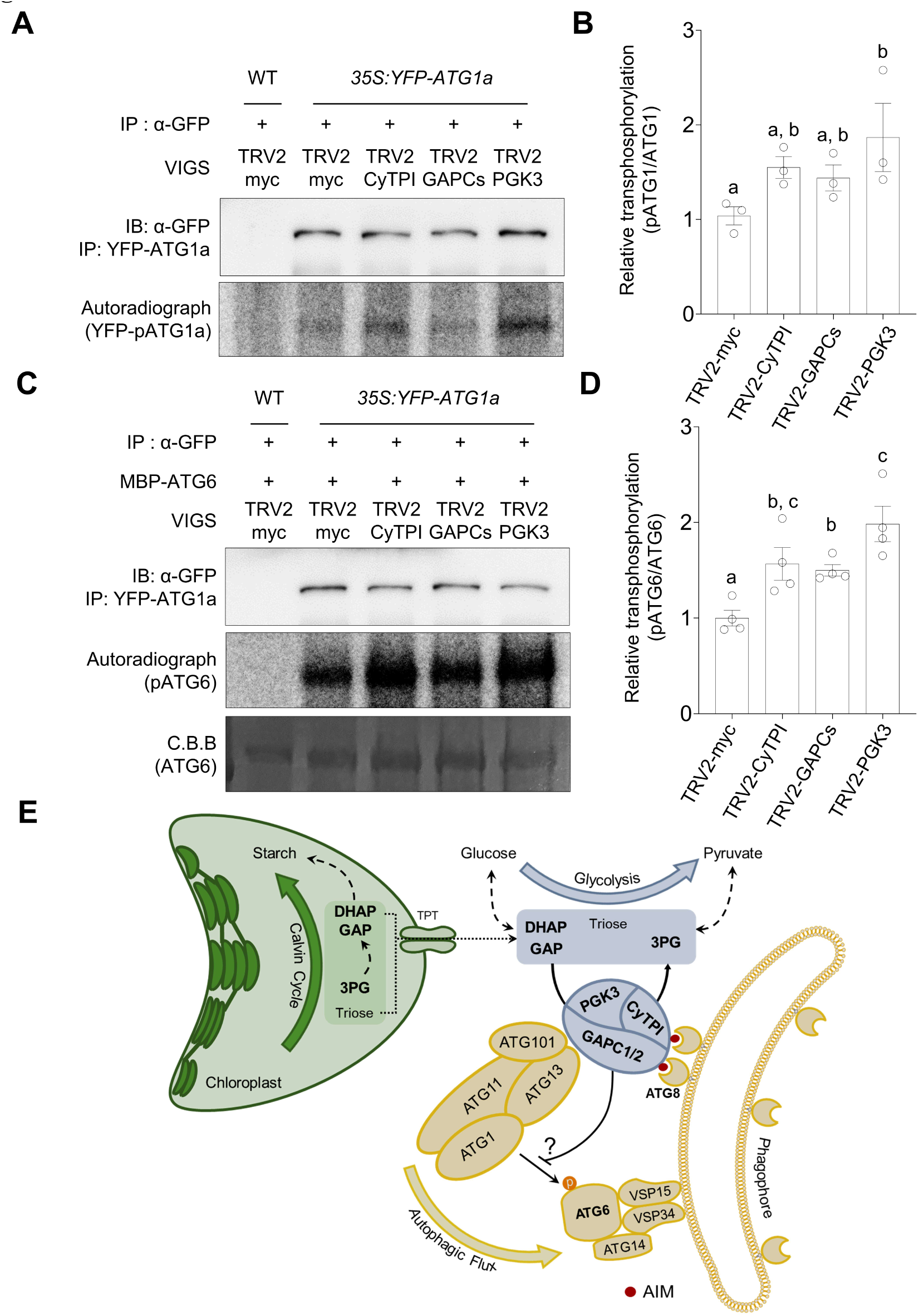
Silencing of the cytosolic TGP enzymes results in increased ATG1 kinase activity. (**A**) Autophosphorylation activity of ATG1a. *In vitro* kinase assay was performed with immunoprecipitated YFP-ATG1a from VIGS plants. The autoradiograph images indicate autophosphorylation levels of YFP-ATG1a in different VIGS plants. Immunoblotting using anti-GFP antibody showed the level of immunoprecipitated YFP-ATG1a in each sample. Three independent experiments yielded similar results and representative images are shown. (**B**) Quantification for relative autophosphorylation levels of YFP-ATG8a shown in (**A**). Autoradiograph band intensities were divided by immunoblotting band intensities of YFP-ATG1a, and the ratio is presented relative to the TRV2-myc control. Statistical significance was calculated by linear mixed effect modelling; letters above the boxes (*a* and *b*) indicate the results of Tukey’s post hoc test. (**C**) Transphosphorylation activity of ATG1a toward ATG6. *In vitro* kinase assay was performed with immunoprecipitated YFP-ATG1a from VIGS plants and purified recombinant MBP-ATG6 protein. The autoradiograph image indicates phosphorylated MBP-ATG6 by YFP-ATG1a. Coomassie brilliant blue (CBB) staining shows input levels of MBP-ATG6 proteins. Four independent experiments yielded similar results and representative images are shown. (**D**) Quantification for relative phosphorylation levels of MBP-ATG6 shown in (**C**). Autoradiograph band intensities were divided by CBB-stained MBP-ATG6 intensities, and the ratio is presented relative to the TRV2-myc control. Statistical significance was calculated by linear mixed effect modelling; letters above the boxes (*a-c*) indicate the results of Tukey’s post hoc test. (**E**) A schematic model of autophagy regulation mechanisms by the cytosolic TGP enzymes. The TGP enzymes (CyTPI, PGK3, and GAPC1/2) in the cytosolic glycolysis pathway, which use triose phosphates transported from chloroplasts via TPT antiporter, play a role in autophagy regulation. The enzymes interact with ATG101 and ATG13 subunits of the ATG1 kinase complex that controls the initiation and growth of the autophagosomes, and suppress an increase in autophagic flux by inhibiting the ATG1 kinase activity.

## Discussion

Nutrient availability is a weighting factor for the balance between catabolism and anabolism (Gonzalez et al., 2020). As most phototrophic organisms mainly produce energy and carbonic materials via photosynthesis, the mechanisms of autophagy regulation via carbon starvation may be different from those of heterotrophic organisms. Recent evidence suggests that plants may regulate autophagy differentially in response to carbon and nitrogen starvation (Huang et al., 2019b). For example, plants induce autophagy upon nitrogen starvation only through ATG1 kinase, but in response to carbon starvation, use both ATG1–dependent and –independent mechanisms (Huang et al., 2019b). These complex regulations indicate that plants may have evolved to manage autophagy more flexibly in response to carbon depletion because of their photoautotrophic nature. Plants accumulate starch during the daytime and consume it at the nighttime. It has been previously proposed that autophagy may also contribute to supplying energy during the nighttime (Izumi et al., 2013). We hypothesized that plants may have specific mechanisms for monitoring carbon availability to modulate autophagic flux.

To address this hypothesis, we performed VIGS screening; subsequent analyses suggested that autophagic flux is modulated by consecutive TGP enzymes (CyTPI, GAPC1/2, and PGK3) in the cytosolic glycolysis pathway (Figure 1A, B). Indeed, carbohydrates produced from the Calvin-Benson cycle in chloroplasts are transported to the cytosol in a triose phosphate form by a triose-phosphate/phosphate translocator (TPT) located in the chloroplast envelope (Walters et al., 2004). Structural analysis revealed that only 3^rd^ carbon-phosphorylated trioses (i.e., DHAP, GAP, and 3-PG) and inorganic phosphate are preferentially transported by the TPT antiporter (Lee et al., 2017b). These triose phosphates, transported from chloroplasts through TPT, can become substrates of TGP enzymes in the cytosol. Moreover, these three enzymes interact with each other in the cytosol, possibly forming a protein complex (Figure 1F; Supplemental Figure 1F). The three TGP enzymes seemed to be located at a strategic point in the cytosolic glycolysis pathway to monitor photosynthetic efficiency by sensing the availability of their substrates (Figure 1B; Supplemental Figure 1D).

In this study, we identified novel interactions between TGP enzymes and the ATG1 regulatory subunits ATG101 and ATG13 (Figure 3A; Supplemental Figure 3A). Considering the upregulated ATG1a kinase activity in the TGP VIGS plants (Figure 7), the cytosolic TGP enzymes can negatively regulate autophagy by repressing the activity of the ATG1 complex and possibly by blocking access to its downstream components. Triose phosphates transported from chloroplasts may play a role in the repression of autophagy by binding to TGP enzymes as substrates, thereby maintaining the active conformation of the enzymes in the cytosol. Therefore, the interactions between cytosolic TGP enzymes and ATG101/ATG13 may connect photosynthetic efficiency to autophagy induction. ATG101 often interacts with CyTPI and GAPCs in large foci structures in an AIM sequence-dependent manner (Figure 3). Interestingly, the AIM sequences in CyTPI and GAPCs were highly conserved in most land plants, including ferns, lycophytes, and bryophytes (Figure 3C). Considering that these land plants also possess all the components of the ATG1 complex, including ATG101, the mechanism of autophagy regulation by TGP enzymes may be conserved in most land plants. Collectively, our results revealed a plant-specific mechanism of autophagy regulation, which employs three consecutive triose phosphate-processing glycolytic enzymes. In addition, when our manuscript was about to be submitted, Guan et al. (2022) reported that GAPCs and PGK3 are involved in the regulation of autophagy by heightening the inhibition of ATG3-ATG8e and ATG6-VPS34 interactions by phosphatidic acid. Thus, TGP enzymes are shown to be involved in the regulation of autophagy, responding to photosynthetic activity and the second messenger molecules, but detailed mechanisms behind these regulations remain to be uncovered.

The mutants of the TGP enzyme genes exhibited sugar-dependent abnormalities in leaf development near shoot apical meristem (SAM) (Figure 2). Activation of SAM relies on both light signaling and photosynthesis-driven nutrient sensing, and both pathways converge at the activation of TOR kinase (Pfeiffer et al., 2016; Li et al., 2017). Considering that the active TOR kinase firmly inhibits nutrient starvation-induced autophagy (Pu et al., 2017), the cellular state of autophagic flux may influence stem cell maintenance/differentiation, depending on nutrient availability. Indeed, a recent study revealed that autophagy contributes to root meristem size regulation in a glucose-dependent manner (Huang et al., 2019b). These findings support that sugar-dependent abnormal leaf formation in TGP mutants may be related to dysregulation of autophagic flux in SAM. Furthermore, cell elongation is regulated by the stability of BRASSINAZOLE RESISTANT 1 (BZR1) and BRI-EMS SUPPRESSOR1 (BES1) transcription factors via selective autophagy (Nolan et al., 2017; Yang et al., 2017). BZR1 and BES1 are ubiquitinated and docked in the autophagic body via DOMINANT SUPPRESSOR OF KAR2 (DSK2), an autophagic adaptor protein (Nolan et al., 2017). Moreover, BZR1/BES1-related selective autophagy is modulated by the TOR pathway, which controls cell elongation according to nutrient availability (Zhang et al., 2016). Accordingly, the cell division/elongation defects of the cytosolic TGP enzyme mutants at low sucrose concentrations (Figure 2; Supplemental Figure 2) may be related to the uncontrolled degradation of unidentified target proteins via autophagy. Indeed, the leaf developmental defect of the TGP mutants was alleviated upon crossing with *atg101-1* (Figure 5). This study is the first to examine the loss-of-function phenotypes of *ATG101* in plants. The *atg101-1* mutant exhibited typical autophagy-deficient phenotypes, similar to other subunits of the ATG1 complex (Figure 4). Taken together, our results suggest that CyTPI, GAPCs, and PGK3 not only regulate autophagy, acting epistatically upstream of the ATG1 complex, but may also coordinate cell division/elongation through autophagy regulation. Identification of further interacting partners of CyTPI, GAPCs, and PGK3 may help explain how plants modulate growth via autophagy according to nutrient availability.

## Materials and methods

### Plant Materials and Growth Conditions

Seeds of *A. thaliana* (Col-0 ecotype) expressing GFP-ATG8a under the control of the cauliflower mosaic virus 35S promoter (35Sp) was obtained from Dr. Taijoon Chung (Pusan University, Korea). T-DNA insertion mutant lines *hxk1-3* (SALK_070739), *cytpi* (GK_244H09), *gapc1-1* (GK_268H09), *gapc2-1* (SALK_016539), *pgk3-1* (SALK_062377), *pgk3-2* (SALK_066422), *atg101-1* (WiscDsLox337F01), *atg5-1* (SAIL_129_B07), and *atg11-1* (SAIL_1166_G10) were obtained from the Arabidosis Biological Resource Center. Genotyping of the mutants was performed by the standard methods using gene-specific primers (Table S1). The *gapc1/2* double mutant was generated by crossing between the *gapc1-1* and *gapc2-1* mutants as previously described (Guo et al., 2012). The *atg13a-1* (SALK_065240) *atg13b-2* (GK_510F06) double mutant (*atg13a/atg13b*) was kindly provided by Dr. Faqiang Li (South China Agricultural University, China). The double mutants *hxk1-3 × atg101-1*, *cytpi × atg101-1, gapc1/2 × atg101-1*, *pgk3-1 × atg101-1*, and *pgk3-2 × atg101-1* were generated by crossing between the mutants, and T3 homozygous seeds were used in this study. Complementation lines for *atg101-1* were generated by floral dipping with *Agrobacterium* C58C1 strain containing the pCAMBIA-ATG101p(-628 bp):YFP-ATG101 construct, and two independent T3 homozygous lines (C13 and C14) were used in this study.

For VIGS and transient expression, *A. thaliana* and *N. benthamiana* plants were grown in soil in a growth chamber (22℃, 60% humidity, 100-120 μmol m^−2^ s^−1^ light intensity using light bulbs [Philips TLD36W/865/FL40SS/36/EX-D], and 16-h-light/8-h-dark cycle). For the liquied culture, *Arabidopsis* seeds were surface-sterilized with 70% ethanol, incubated at 4℃ for 1 day for stratification, and then sown in 6-well or 12-well plates containing 1 mL or 0.5 mL of liquid medium (1/2 MS basal salt medium [Duchefa, M0221], pH 5.7 adjusted with KOH), respectively. After germination, seedlings were grown in 1/2 MS liquid medium containing 30 mM glucose, which was changed every other day. For the analysis of growth phenotype, *Arabidopsis* seeds were surface-sterilized and imbibed, and then grown in plates (1/2 MS basal salts, pH 5.7 adjusted by MES-KOH, and 0.6% phytoagar [Duchefa, P1001]) containing various sugar concentrations.

### Virus induced gene silencing (VIGS)

VIGS was performed in *Arabidopsis* as described previously (Lee et al., 2017a), using soil-grown seedlings at two to four true-leaf stages. The cDNA fragments of *Arabidopsis* TOR complex genes (*TOR, LST8-1* and *RAPTOR1A/1B*) and forty five metabolic genes were cloned into pTRV2 vector (Burch-Smith et al., 2006) using In-Fusion cloning (Takara Korea). Both pTRV1 (contaning a gene encoding viral RNA-depenent RNA polymerase) and pTRV2 (contaning a cDNA fragment of a target gene) were introduced into *Agrobacterium* (GV3101 strain). pTRV2-Myc was used as a negative control for VIGS. The transformated *Agrobacterium* strains were cultured overnight in Luria-Bertani medium containing 10 mM MES-KOH (pH 5.7), 20 μM acetosyringone, kanamycin (50 μg/mL), and rifampicin (50 μg/mL). The *Agrobacterium* culture was harversted and resuspended at OD_600_ of 1.5 in the infiltration medium (10 mM MgCl_2_, 10 mM MES-KOH, pH 5.7, and 200 μM acetosyringone). The mixture was then incubated at 22℃ for more than 4 h and infiltrated into the 1^st^ true leaf of *Arabidopsis* seedlings using a needless syringe.

### Phagosome visualization

Leaves from VIGS plants were observed by fluorescence microscope (Olympus, BX51) and confocal laser scanning microscope (Carl Zeiss, LSM 880) for initial screening and Z-stacked image processing (Maximum Intensity Z-projection), respectively. Puncta numbers were counted by Fiji software using the “Analyze Particles” method.

### BiFC

BiFC was performed using the muticolor bimolecular fluorescence complementation system, as previously described (Gehl et al., 2009). Coding sequences (CDSs) of *HXK1, CyTPI, GAPC1, GAPC2, PGK3, ATG101, ATG13a* and *ATG13b* were cloned into the Fu79 entry vector (Wang et al., 2013), and *PdTPI, PGK2* and *PGK3* were cloned into the pENTR/D-TOPO entry vector (Thermo Fisher Sientific). These entry vectors were used for subcloning into the destination vectors pDEST-^GW^VYNE and pDEST-^GW^VYCE for Fu79 system, and pDEST-VYNE(R)^GW^ and pDEST-VYCE(R)^GW^ for pENTR/D-TOPO system, using LR clonase (Thermo Fisher Scientific, 11791100). In addition, *CyTPI, GAPC1* and *PGK3* entry clones were used for subcloning into the pDEST-^GW^SCYNE and pDEST-^GW^SCYNE vectors.

To analyze the AIM motif mutant, we performed target-directed mutagenesis based on In-Fusion cloning. CDSs of *CyTPI* and *GAPC1* containing the AIM motif were cloned into Fu79 entry vector and then subcloned into the pDEST-^GW^VYNE and pDEST-^GW^VYCE vectors. *Agrobacterium GV3101* strains containing the VYNE (and/or SCYNE), VYCE (and/or SCYCE), and 35Sp:p19 construct were coinfiltrated at the adjusted OD_600_ as 0.7:0.7:1 into leaves of 3-week-old *N.bethamiana* plants. BiFC signals were monitored 36∼48 h after infiltration in the abaxial side of leaf epidermis using a confocal laser scanning microscope (Carl Zeiss, LSM 700). To combine colocalization and BiFC, *Agrobacterium GV3101* strains containing the SCYNE, SCYCE, pCAMBIA1390-35Sp:mCherry-ATG8a and 35S:p19 construct were coinfiltrated at the adjusted at the OD_600_ ratio of 0.7:0.7:0.2:1 into leaves of *N. benthamiana* plants. In this case, fluorescence was monitored in protoplasts 36 h after infiltration. To detect protein expression, immunoblotting was performed with 50 μg of protein extracts using anti-Flag-M2-HRP (Sigma-Aldrich, A8592; 1:10,000), anti-c-myc-peoxidase (Roche, 11814150001; 1:10,000) and anti-HA-peroxidase (Roche, 12013819001; 1:10,000) conjugated antibodies for SCYNE, VYNE and VYCE (or SCYCE), respectively.

### Quantitative RT-PCR

Total RNA was extracted using the IQeasy Plus Plant RNA Extraction Mini kit (iNtRON Biotechnology; Korea) according to the manufacturer’s instructions. cDNA synthesis was performed with one μg of total RNA using the RevertAid First Standard cDNA synthesis kit (Thermo Fisher Scientific, K1622) and oligo(dT) primers, according to the manufacturer’s instructions. Real-time qPCR was performed in a 96-well plate using diluted cDNAs (1:100), RealHelix^TM^ qPCR kit (NANOHELIX; Korea), and the StepOnePlus Realtime PCR System (Applied Biosystems, 4376600) with specific primer sets (Table S1). Normalization was performed as described in the figure legends, using *PP2AA3* (protein phosphatase 2A subunit A3) mRNA as a reference gene.

### Carbon starvation treatment

Carbon starvation was performed as previously described by as “re-greening assay” with minor modification (Lee et al., 2017a). Briefly, twelve-day-old seedlings in liquid culture were washed 4 times with the 1/2 MS medium without carbon source, and then incubated in the dark for 5 days in fresh 1/2 MS medium without carbon source, designated as “dark/starvation (DS)” treatment. After DS, the seedlings were supplied with fresh 1/2 MS medium containing 30 mM glucose and incubated in the light for 3 days or 5 days.

### Nitrogen starvation treatment

Seedlings were grown for 10 days in MS medium (pH 5.7 adjusted by MES-KOH) with 1% sucrose and 0.8% phytoagar, and then transferred to MS medium that contains or lacks nitrogen source for 5 days.

### Chlrophyll mesurement

Chlorophyll contents were measured as described (Lee et al., 2017a). Chlorophyll was extracted from seedlings using aqueous 80% acetone. Absorbance of the extract was monitored at 663.6 and 646.6 nm wavelength by VersaMax Absorbance Microplate Reader (Molecular Devices). The total chlorophyll contents were calculated by the equation as previoulsy described (Porra and Scheer, 2000), and normalized by plant fresh weight.

### Immunoblotting

To measure autophagic flux, immunoblotting was performed with twelve-day-old seedlings grown in liquid culture. Homogenized samples were resuspended in the same volume of modified extraction buffer (50 mM Tris-HCl pH 7.5, 150 mM NaCl, 1% NP-40, 0.1% SDS, 0.5% sodium deoxycholate, 2 mM Na_3_VO_4_, 2 mM NaF, 20 mM β-glycerophosphate, and cOmplete protease inhibitor cocktail [Roche, 11836170001]). After centrifugation, NuPAGE^TM^ LDS sample buffer (Thermo Fisher Sientific, NP0007) was added for incubation in 70℃ for 10 min. Twenty five μg of total protein was subjected to SDS-PAGE (4-20% in Tris-glycine buffer) using low voltage (100 V) for 2.5 h at 4℃. Immunoblotting was performed with the rabbit polyclonal anti-ATG8 antibody (Agrisera, AS14 2769; 1:5,000 in 5% BSA), anti-GAPC1/2 (Agrisera, AS15 2894; 1:10,000 in 5% skimmed milk), anti-PGK3 (produced from rabbits by repeated injection using recombinant His-PGK3 protein; 1:10,000 in 5% skimmed milk, GW Vitek, Korea), and mouse monoclonal anti-GFP antibody (Roche, 11814460001; 1:5,000 in 5% skimmed milk). Signals were detected carefully to avoid band saturation by Imagequant LAS 4000 (GE Healthcare Life Sciences). Ponceau S staining (Sigma-Aldrich) was performed after signal detection.

### Protoplast isolation and transfection

*Arabidopsis* mesophyll protoplasts were isolated as described previously (Yoo et al., 2007). Briefly, leaves from 4-week-old *Arabidopsis* plants (grown in 16 h light/8 h dark) were cut into strips and then incubated in the digestion solution (20 mM MES-KOH pH 5.7, 1.5% Celluase R-10 [Yakult; Japan], 0.4% Macerozyme R-10 [Yakult; Japan], 0.4 M mannitol, 20 mM KCl, 10 mM CaCl_2_, and 0.1% BSA) for 3 h in the dark. The protoplasts were washed using W5 buffer (2 mM MES-KOH pH 5.7, 154 mM NaCl, 125 mM CaCl_2_, and 5 mM KCl) and collected by centrifucation at 100 *g* for 2 min. For autophagosome observation, protoplasts (10^5^ cells) were transfected with 10 μg of p326-sGFP-ATG8a plasmid DNA for 16 h in WI solution (4 mM MES-KOH pH 5.7, 0.5 M mannitol, and 20 mM KCl). For Concanmycin A (APExBio, A8633) treatment, 1 μM Con A was treated for 16 h along with transfection.

### *In vitro* kinase assay using immunoprecipitated ATG1a

To purify substrate of ATG1a, CDS of ATG6 was cloned into the pMal-C2X vector (New England Biolabs) for MBP fusion using infusion cloning (Takara Korea). The constructrs were transfromed into *E.coli* BL21 (DE3) strain. Cells were grown in LB medium containing 1% glucose and ampicillin (50 μg/mL) at 37℃ to an OD_600_ of 0.4, and induced by 0.25 mM IPTG for 16 h. MBP-fusion proteins were purified using Amylose Resin (New England Biolabs, E8021), following the manufacturer’s instruction with minor modification. During purification, we used a single buffer (20 mM Tris-HCl and 200 mM NaCl) and added 10 mM maltose into the buffer for protein elution. After purification, proteins were concentrated using Amicon Ultracel 30K (Merck, UFC503096) according to the manufacturer’s instruction.

Kinase assay was performed as previously described with minor modifications (Sanchez-Wandelmer et al., 2017). YFP-ATG1a protein was expressed in *N. benthamiana* leaves or *Arabidopsis* stable transgenic line. Samples were ground on liquid nitrogen and mixed with IP kinase assay buffer A (100 mM Tris-HCl, pH 7.5, 300 mM NaCl, 10 mM EDTA, 10 mM EGTA, 1% NP-40, 10 mM Na_3_VO_4_, 10 mM NaF, 1 mM phenylmethylsulfonylfluoride, 10% glycerol, and protease inhibitor cocktail [Roche, 11836170001]). After brief centrifugation, GFP-Trap agarose resin (Chromotek, gta-10) was added into the supernatant and incubated for 4 h at 4℃ with rotation. After incubation, the agargose resin was wahsed 4 times with IP kinase assay buffer A without glycerol, followed by washing twice with IP kinase assay buffer B (20 mM MOPS, pH 7.5, 1 mM EGTA, 10 mM Na_3_VO_4_, and 15 mM MgCl_2_). Kinase assay was performed with 2 μCi [γ-^32^P]-ATP, immunoprecipitated YFP-ATG1a, and 5 μg recombinant substrate proteins in 20 μL IP kinase buffer B for 20 min at 30℃. The reaction was stopped by adding Laemmli sample buffer and incubated for 5 min. After SDS-PAGE, the gel was stained with Sun-Gel staining solution (LPS Solution; Korea) and dried using a gel dryer (Hoefer^TM^ GD2000 Gel Dryer System; Fisher Scientific). Radioactivity was deteced using a phosphorimager (BAS-2500, Fujifilm). Immunoblotting of immunoprecipitated YFP-ATG1a was performed with the anti-GFP antibody (Roche, 11814460001).

### Statistical analysis

GraphPad PRISM 9.0 and R & Bioconductor (Ver. 4.1.2) were used for most data analyses and visualization. Outliers were determined and removed using the ROUT test, performed in GraphPad PRISM. Two-sides unpaired t-test and One-way anlysis of variance (ANOVA) were implemented in GraphPad PRISM and R & Bioconductor. Post hoc tests following analysis of variance were specified in the figure legends.

## Author Contributions

D.-H.L. and H.-S.P. conceived, designed, and coordinated the project. D.-H.L. and I.C. performed all the experimental works with S.J.P. and S.K. D.-H.L., I.C., M.-S.C., H.-S.L. and H.-S.P. analyzed the results. D.-H.L. and H.-S.P. wrote the manuscript. All authors discussed the results and commended on the manuscript.

## Acknowledgement

We thank Dr. Taijoon Chung (Pusan National University) and Tae-Wuk Kim (Hanyang University) for helpful discussions. We are grateful to Dr. Faqiang Li (South China Agricultural University), Dr. Taijoon Chung, and Dr. Richard D. Vierstra (Wisconsin University) for generously providing the seeds of *35Sp:GFP-ATG8a* and *atg13a-1/atg13b-2*.

This research was supported by the Basic Science Research Program (2018R1A6A1A03025607) and the Mid-Career Researcher Program (2022R1A2C1009088) from the National Research Foundation (NRF) of the Republic of Korea. D.-H. Lee was supported by BK21 Fellowship of Yonsei University.

## Supplemental Materials

**Supplemental Figure 1.**
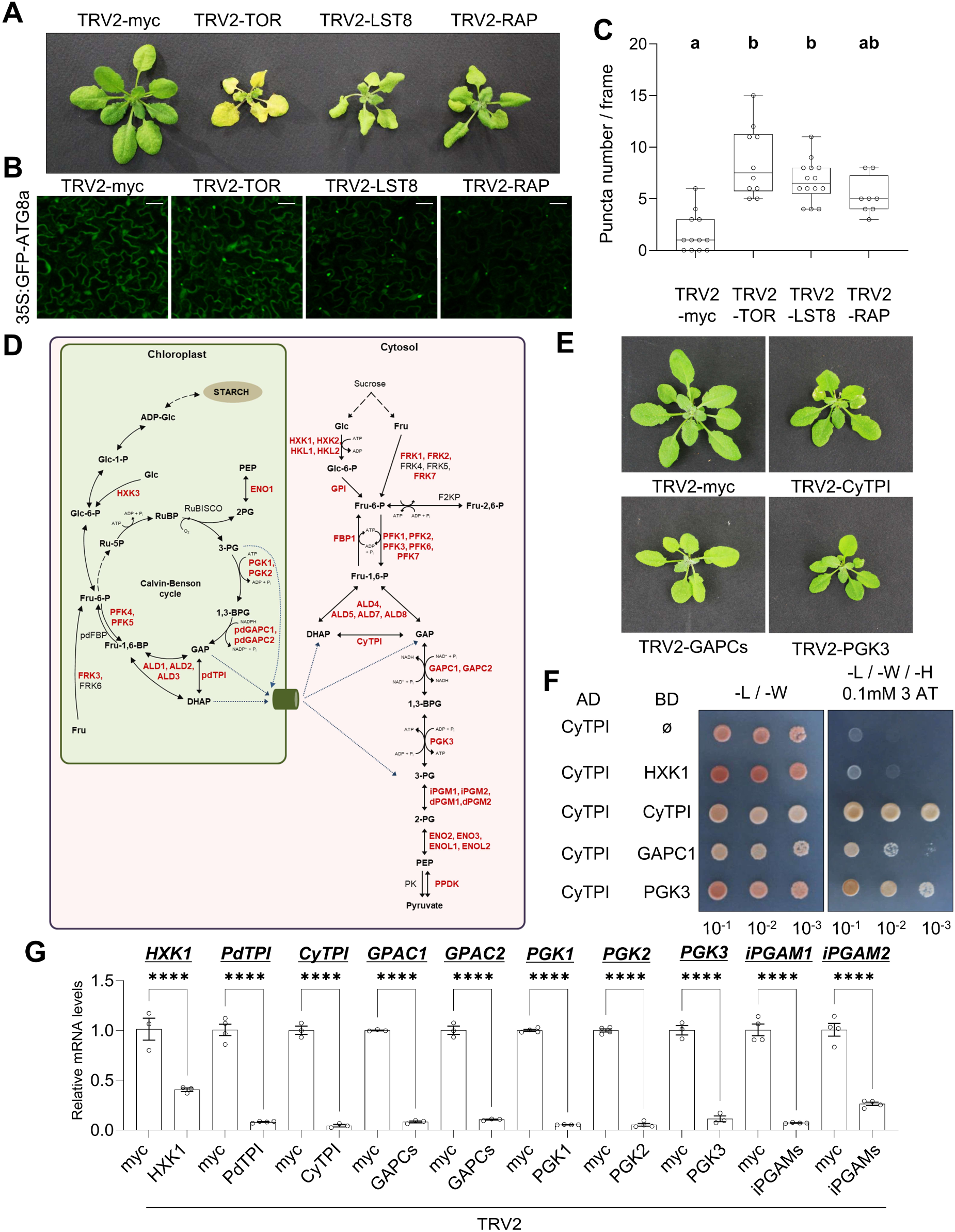
VIGS screening of cytosolic and chloroplast enzyme genes that are related with glycolysis and Calvin-Benson cycle. (**A**) VIGS phenotypes of TOR complex (TORC) genes, *TOR, LST8*, and *RAPTOR*, in Arabidopsis. TRV2-myc was used as the control for VIGS. Photos were taken 14 days after infiltration (DAI). (**B**) Confocal microscopy of GFP-ATG8a fluorescence (Z-stack image projection) in epidermal cells of the *Arabidopsis* leaves after VIGS of TORC genes at 14 DAI. Scale bar = 10 μm. (**C**) Puncta numbers per frame of confocal microscopy images (Z-stack image projection). The box plot represents the 1^st^ and 3^rd^ quartiles, and whiskers include the min and max of the data points. Dots indicate individual observations. Letters above the boxes (*a* and *b*) indicate the results of one-way ANOVA followed by Tukey’s multiple comparison test (p < 0.05). (**D**) Schematic representation of the glycolysis- and Calvin-Benson cycle-related enzymes, which were selected for VIGS screening. Enzymes indicated in *red* are the targets for VIGS screening. Abbreviations are as follows: Glc, glucose; Fru, fructose; Glc-6-P, glucose-6-phosphate; Fru-6-p, fructose-6-phosphate; Fru-2,6-p, fructose-2,6-bisphosphate; Fru-1,6-p, fructose-1,6-bisphosphate; DHAP, dihydroxyacetone phosphate; GAP, glyceraldehyde 3-phosphate; 1,3-BPG, 1,3-bis-phosphoglycerate; 3-PG, 3-phosphoglycerate; 2-PG, 2-phosphoglycerate; PEP, phosphoenolpyruvate; Ru-5P, ribulose 5-phosphate; RuBP, ribulose 1,5-bisphosphate; Glc-1-P, glucose-1-phosphate; ADP-Glc, ADP-glucose; HXK, hxokinase; GPI, glucose-6-phosphate isomerase; FRK, Fructokinase; FBP, fructose-2,6-bisphosphatase; PFK, phosphofructokinase; ALD, aldolase; TPI, triose phosphate isomerase; GAPC, glyceraldehyde 3-phosphate dehydrogenase; iPGAM, metal ion independent 3-phosphoglycerate mutase; dPGAM, metal ion dependent 3-phosphoglycerate mutase; ENO, enolase; ENOL, enolase like protein; PPDK, pyruvate phosphate dikinase; PK, pyruvate kinase; Rubisco, ribulose 1,5-bisphosphate carboxylase/oxygenase. “Pd” and “Cy” in front of the enzyme name denote the plastidial and cytosolic isoform, respectively. (**E**) VIGS phenotype of the cytosolic TGP enzymes. Photos were taken at 18 DAI. (**F**) Yeast two-hybrid assays for interactions of CyTPI with CyTPI, GAPC1, and PGK3. The BD- and AD-fusion constructs were co-transformed into the yeast strain Y2H Gold. The transformants were grown for 3 days on –L/–W (lacking leucine and tryptophan) control plates (*left panel*) or grown for 5 days on –L/–W/–H (lacking leucine, tryptophan, and histidine) selection plates containing 0.1 mM 3-amino-1,2,4-trizole (3-AT) (*right panel*). BD-ø (empty vector) and BD-HXK1 were used as negative controls. (**G**) qRT-PCR analyses for mRNA levels of selected genes after VIGS. Error bars represent SE from biological replications (n = 3 or 4). Transcripts levels are normalized by *PP2AA3* mRNA, and then expressed relative to those in TRV2-myc. Statistical significance was evaluated using one-way ANOVA followed by Tukey’s multiple comparison test, which represents significance in differences between corresponding experimental data (*, p < 0.05; **, p < 0.01; ****, p < 0.0001).

**Supplemental Figure 2.**
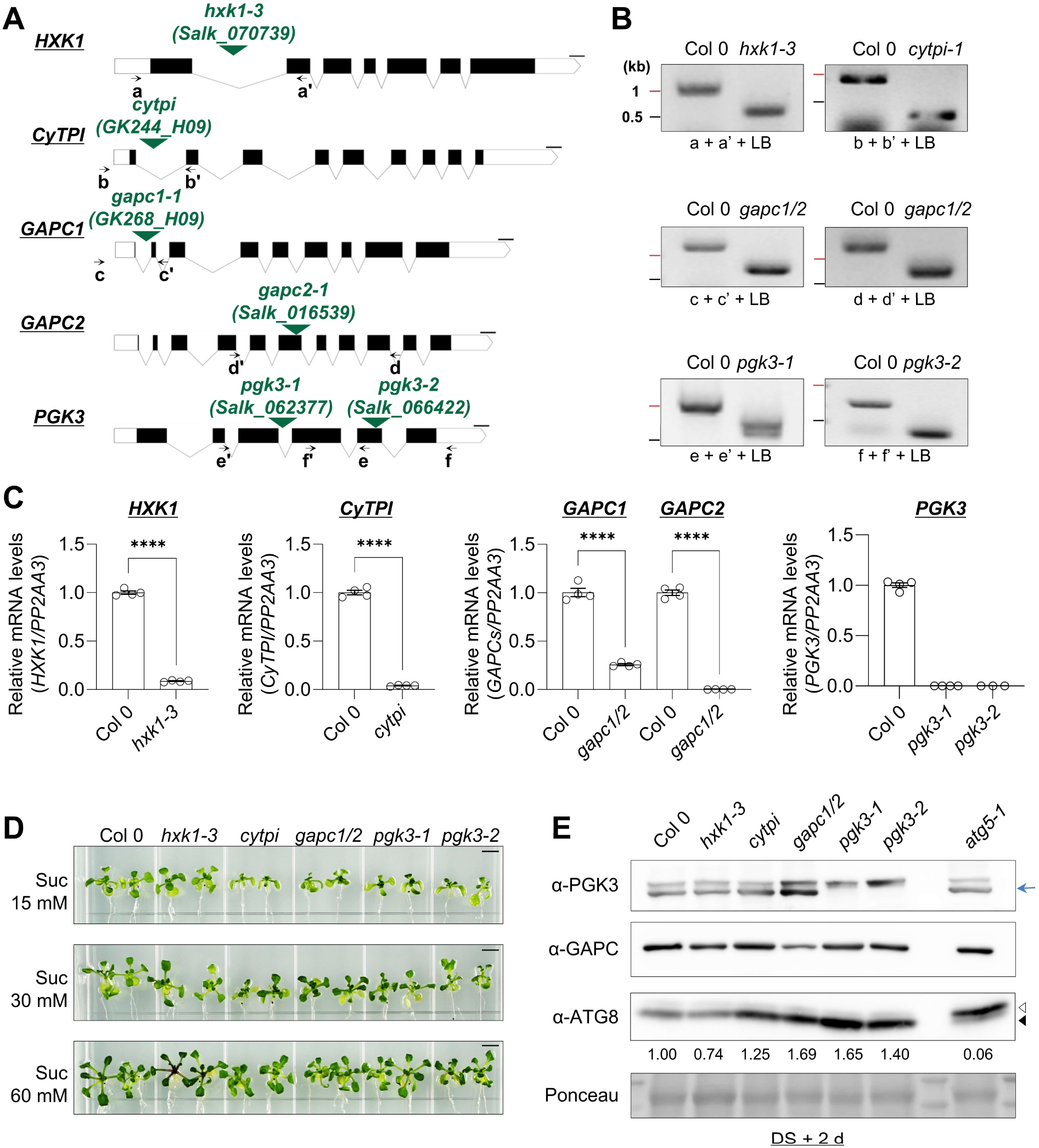
Analyses of the T-DNA mutants of the glycolysis enzymes. (**A**) A schematic presentation of T-DNA insertion sites in the glycolytic enzyme genes. *Green arrowheads* indicate T-DNA insertion sites. *Black arrows* indicate the location and orientation of each primer for genotyping. Scale bars = 100 bp. (**B**) PCR-based genotyping of the glycolysis mutants, *hxk1-3, cytpi, gapc1/2, pgk3-1*, and *pgk3-2*. Three primers used for genotyping of each gene are shown below the PCR data. The PCR data suggested that these mutants are all homozygotes. (**C**) qRT-PCR to determine gene expression level in each mutant. Primer combinations are as described in Supplemental Table 1. Transcript levels are normalized by *PP2AA3* mRNA, and expressed relative to those in Col 0. Error bars represent SE from biological replications (n = 4). Statistical significance was assessed using the unpaired t-test between WT (Col 0) and each mutant. qRT-PCR analyses revealed that *hxk1-3* is a knockdown (KD) allele, while *cytpi, pgk3-1* and *pgk3-2* are knockout (KO) alleles with extremely low transcript levels, in accordance with the previous reports [13,50]. In addition, we generated *GAPC1* and *GAPC2* double mutants using *gapc1-1* (KD) and *gapc2-1* (KO) mutants as reported by Guo et al. (2012), and designated it as *gapc1/2* hereafter. The *gapc1/2* double mutants had about 27% of WT levels of GAPC1 transcripts and undetectable levels of *GAPC2* transcripts. (**D**) Growth phenotypes of the T-DNA mutants in different sucrose concentrations (15, 30, and 60 mM). Photos were taken 20 days after germination. Scale bars = 2.5 mm. (**E**) Immunoblotting for monitoring ATG8-PE:ATG8 patterns in the glycolysis mutants. Twelve-day-old seedlings were incubated in the dark in the medium lacking glucose for 2 days (DS + 2 d). Protein extracts were subjected to immunoblotting with anti-PGK3, anti-GAPC, and anti-ATG8 antibodies. The ATG8 and ATG8-PE are marked with *open* and *closed arrowheads*, respectively. The PGK3 protein band is marked by the blue arrow. Relative band intensities of ATG8-PE/ATG8 are indicated below the anti-ATG8 image.

**Supplemental Figure 3.**
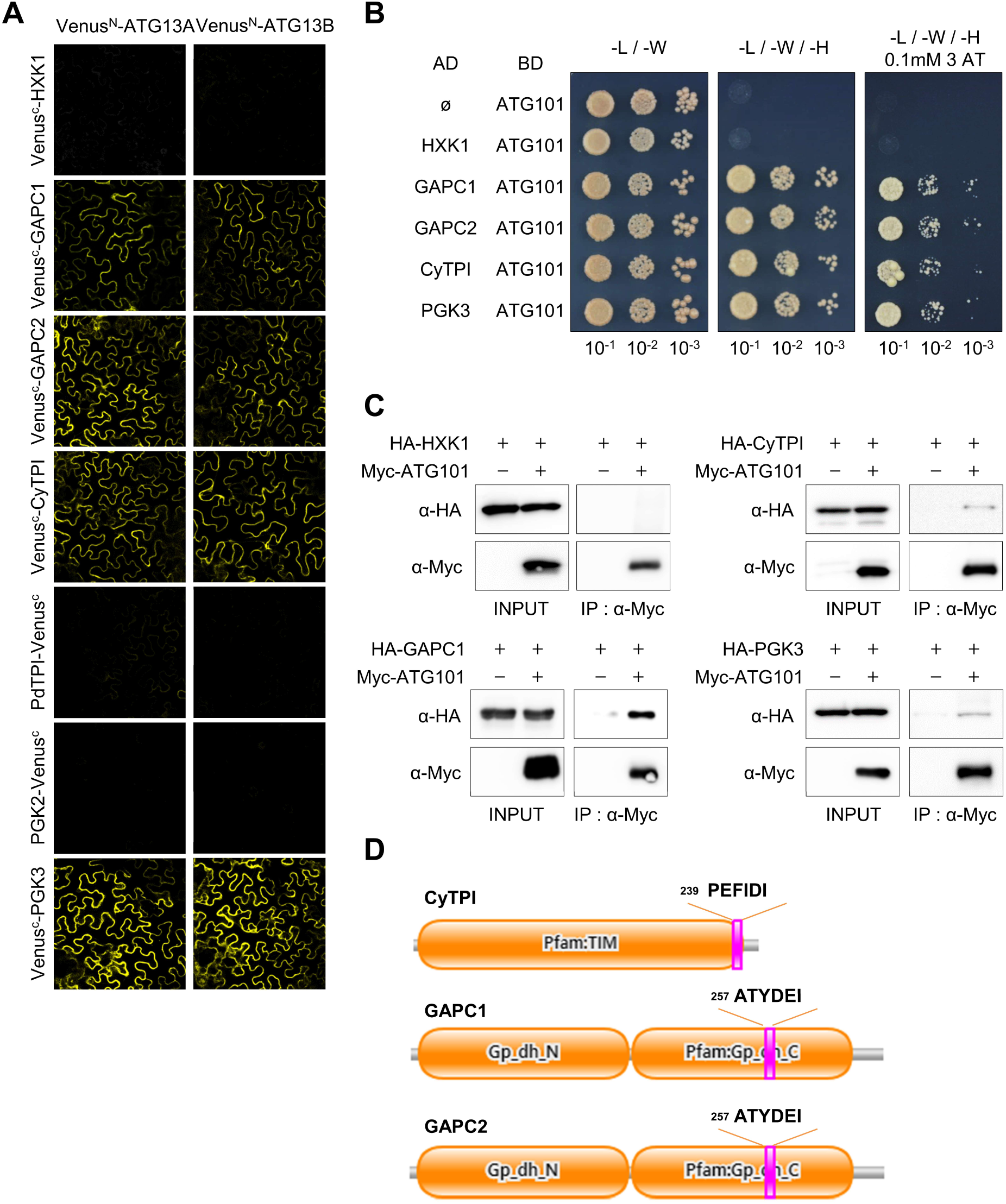
The cytosolic TGP enzymes interact with ATG13 and ATG101. (**A**) BiFC analyses for glycolytic enzymes interactions with ATG13a and ATG13b. VENUS^N^- and VENUS^C^-fused proteins were co-expressed in *N. benthamaia* leaves via agroinfiltration. Leaf epidermal cells were observed by confocal microscopy. Scale bar = 20 μm. (**B**) Yeast two hybrid assay. The glycolytic enzymes were fused to the GAL4 BD domain and co-expressed with GAL4 AD-fused ATG101 in Y2H Gold yeast strain. Serial dilutions of yeast culture were spotted on the diverse medium as indicated. Positive yeast growth on –L/–W/–H medium (lacking leucine, tryptophan, and histidine) in the presence of 0.1 mM 3-AT indicates strong interactions between the bait and prey proteins. (**C**) Co-immunoprecipitation. HA-fused glycolytic enzymes were expressed alone or together with Myc-fused ATG101 in *N. benthamiana* leaves. Total leaf protein extracts were immunoprecipitated with anti-Myc antibody-conjugated resin, and the co-imunoprecipitates were detected by anti-HA antibody. (**D**) The presence of the ATG8-Interacting Motif (AIM) in CyTPI, GAPC1, and GAPC2. Consensus sequences of the AIM (F/Y/W-X_1_-X_2_-L/I/V) were predicted by iLIR program (*in silico* identification of functional LC3-Interacting Region motif; http://repeat.biol.ucy.ac.cy/iLIR/). The predicted AIM sequences and their localization (*pink boxes*) are shown in CyTPI, GAPC1, and GAPC2. Numbers indicate the starting amino acid residue numbers of each sequence.

**Supplemental Figure 4.**
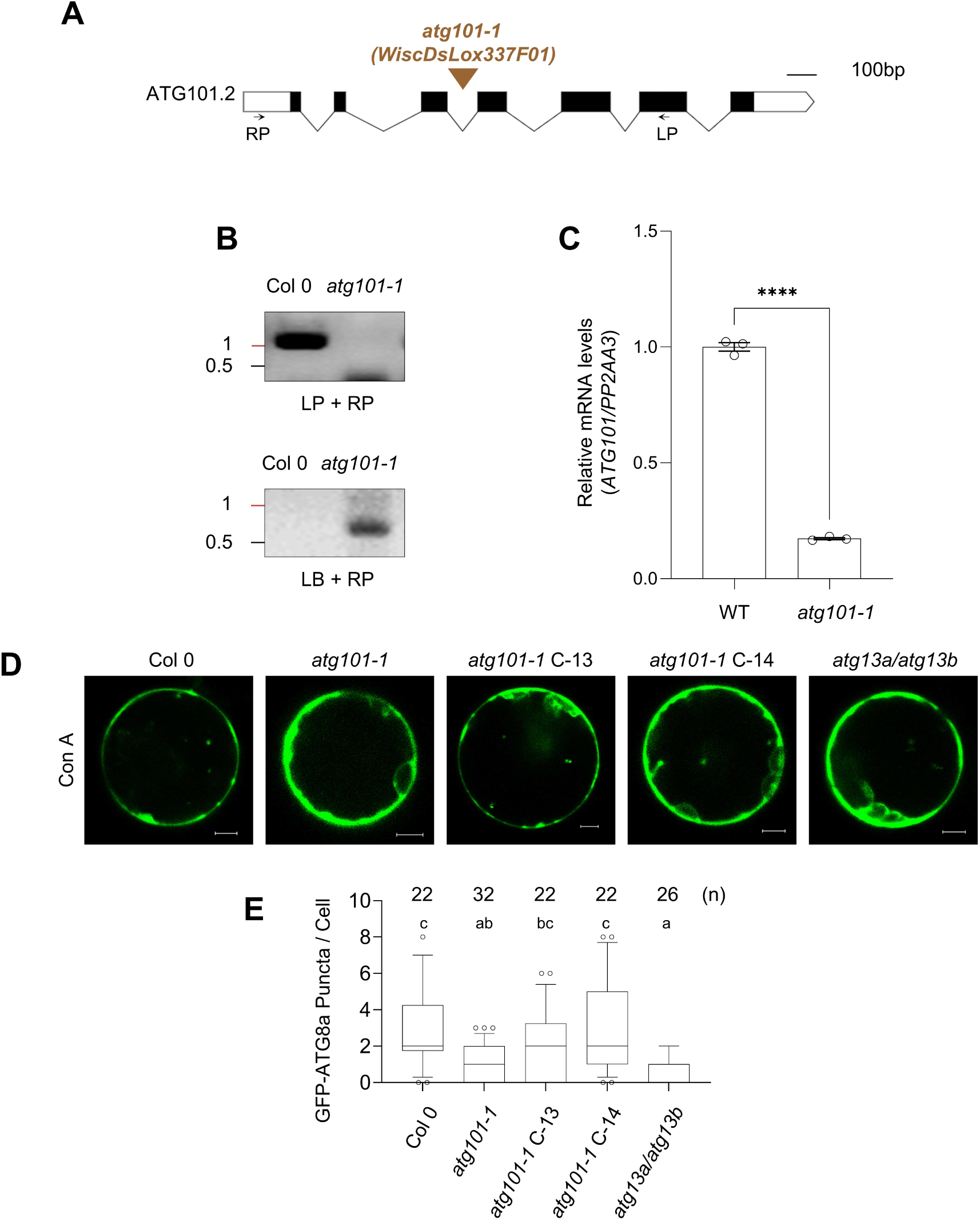
Genotyping for *atg101-1* mutant and its complementation lines. (**A**) A schematic presentation of a T-DNA insertion site in *ATG101*. The *brown arrowhead* indicates the T-DNA insertion site. *Black arrows* indicate the location and orientation of each primer for genotyping. Scale bar = 100 bp. (**B**) PCR-based genotyping of the *atg101-1* mutant. Primer sets are shown under the PCR data. The result suggests that the *atg101-1* mutant is a homozygote. (**C**) qRT-PCR to determine *ATG101* transcript levels in the mutant using primers as described in Supplemental Table 2. Transcripts levels are normalized by *PP2AA3* mRNA, and expressed relative to those in Col 0. Error bars represent SE from three biological replications. Statistical significance was assessed using the unpaired t-test between Col 0 and *atg101-1*. (**D**) Confocal microscopy of GFP-ATG8a fluorescence (Z-stack image projection) in *Arabidopsis* mesophyll protoplasts of the mutants and complementation lines. Leaf protoplasts were transfected with the GFP-ATG8a construct, followed by treatment with 1 μM concanamycin A (Con A). Then GFP-ATG8a puncta in the protoplasts were observed under a confocal microscope. Scale bars = 5 μm. (**E**) The GFP-ATG8a puncta in the protoplasts shown in (**D**) were counted in the Z-stacked confocal microscopy images after Con A treatment. The box plot represents the 1^st^ and 3^rd^ quartiles, and whiskers include the 10^th^-90^th^ percentile of the data points. Letters above the boxes (*a-c*) indicate the results of one-way ANOVA followed by Tukey’s multiple comparison test (p < 0.05).

**Supplemental Figure 5.**
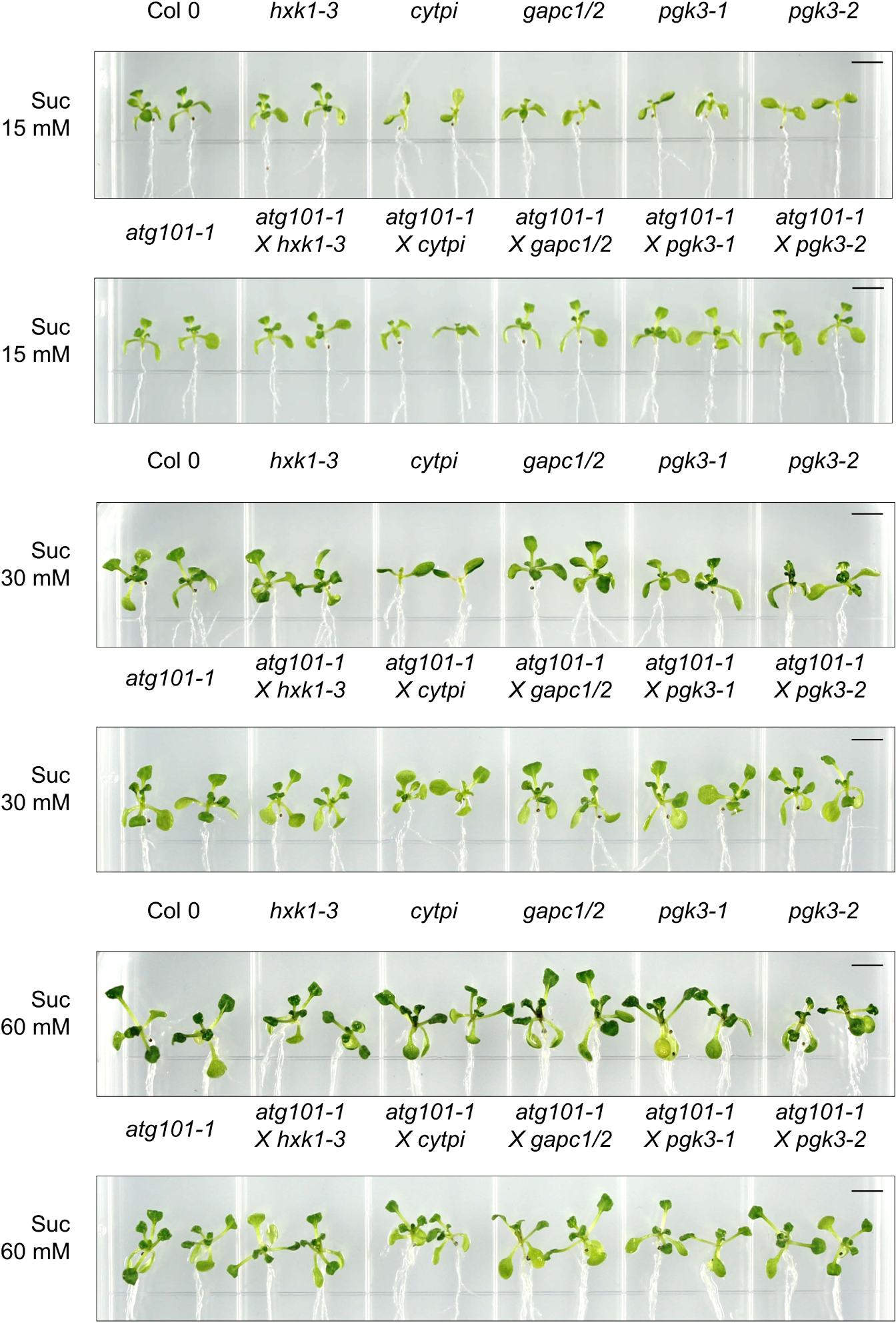
Abnormal phenotypes of the glycolysis mutants were alleviated by crossing with *atg101-1* mutant. Phenotypes of the single T-DNA mutants of the glycolytic enzyme genes and their *atg101*-crossed lines under 15 mM, 30 mM and 60 mM sucrose. Photos were taken 20 days after germination. Scale bar = 2.5 mm.

**Supplemental Figure 6.**
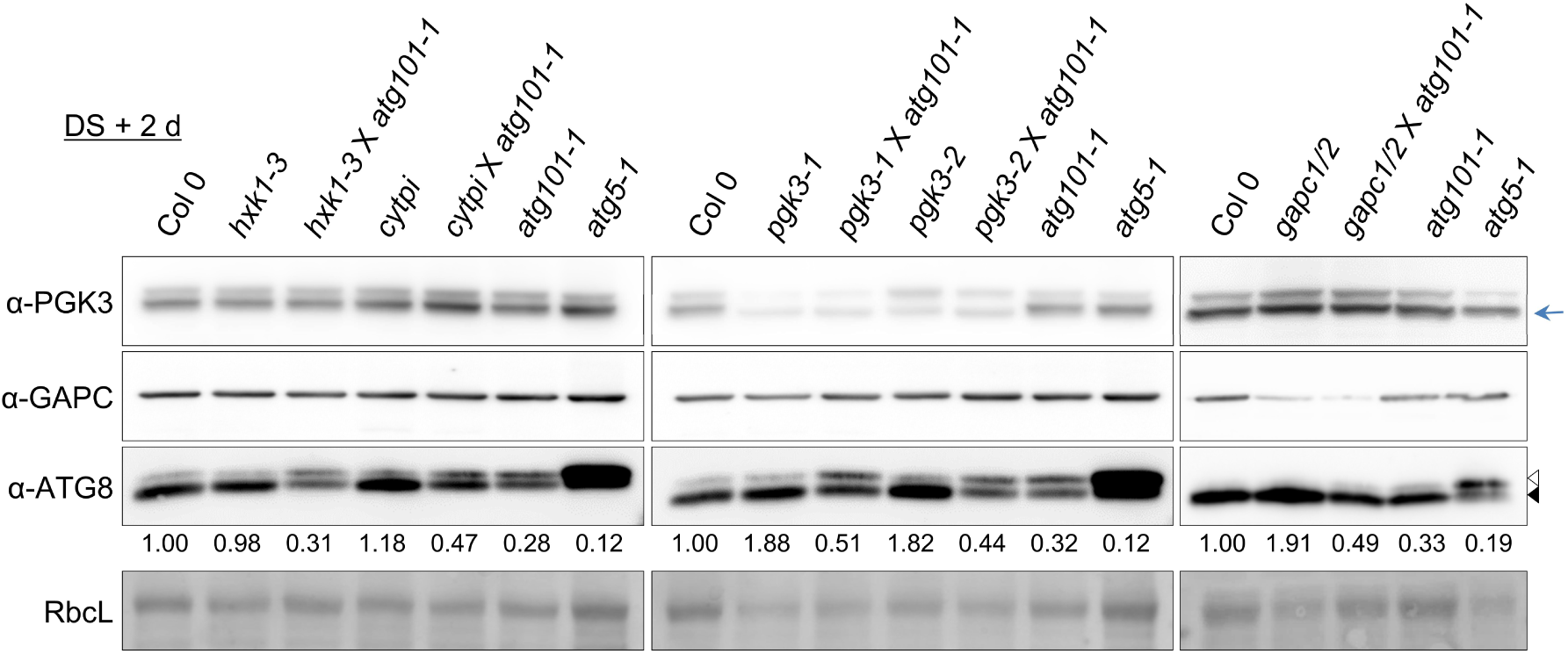
ATG8-PE:ATG8 patterns in the single glycolysis mutants and their *atg101*-cross lines under dark and starvation conditions. Twelve-day-old seedlings were transferred to fresh liquid medium lacking glucose and incubated in the dark for 2 days (DS + 2 d). Then immunoblotting was performed with anti-PGK3, anti-GAPC1, and anti-ATG8 antibodies. The ATG8 and ATG8-PE are marked with *open* and *closed arrowheads*, respectively. The PGK3 protein band is marked by the *blue arrow*. Relative band intensities of ATG8-PE/ATG8 are indicated below the anti-ATG8 image.

**Supplemental Figure 7.**
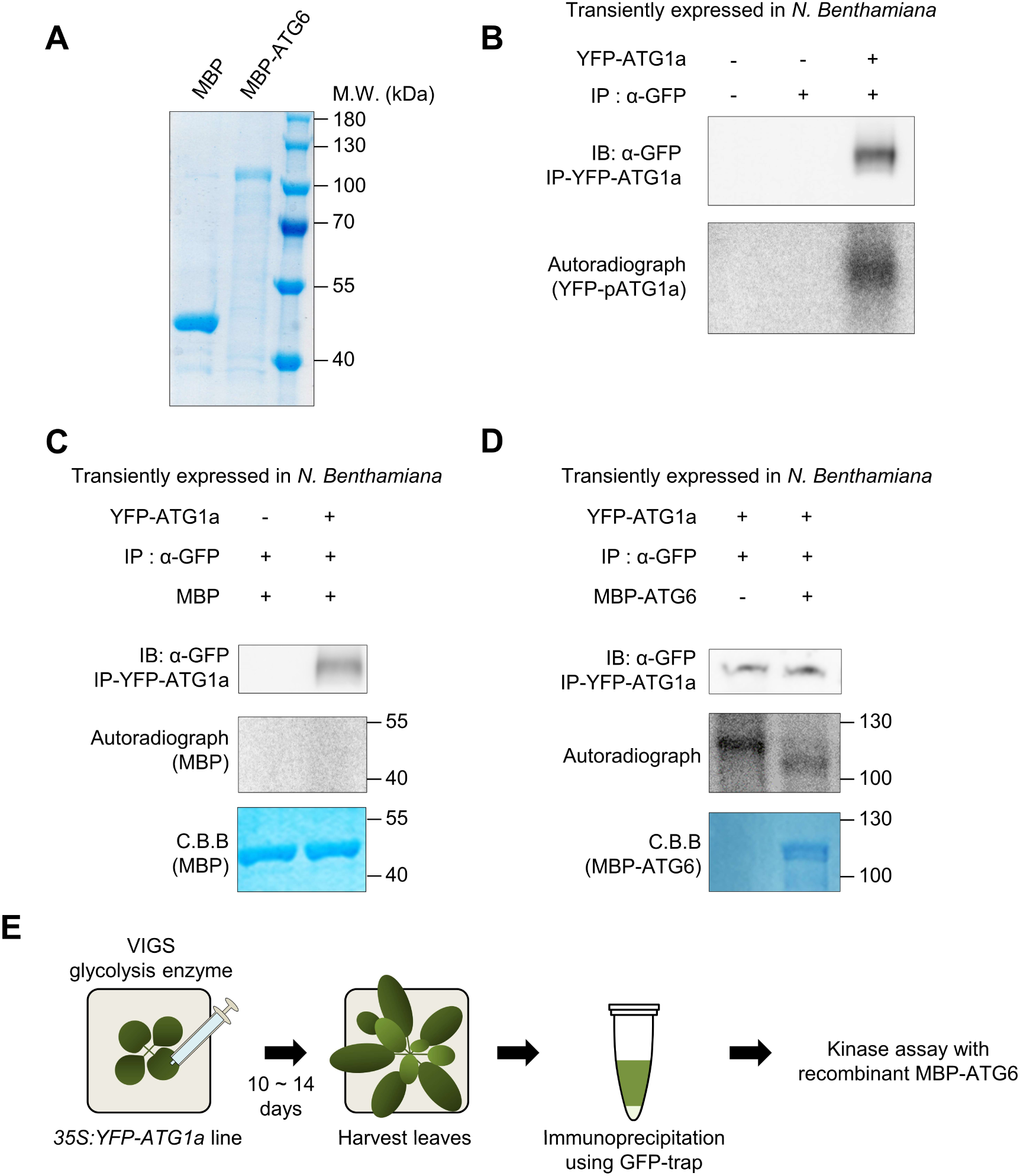
ATG1a kinase phosphorylates ATG6 *in vitro*. (**A**) Purified MBP and MBP-ATG6 proteins. 5 µg of MBP and MBP-ATG6 were subjected to SDS-PAGE and stained by Sun-gel staining solution. (**B**) Autophosphorylation activity of ATG1a. YFP-ATG1a was expressed in *N. benthamiana* leaves via agroinfiltration. Then *in vitro* kinase assay was performed with YFP-ATG1a protein immunoprecipitated from the leaf extract. The autoradiograph image indicates autophosphorylation levels of YFP-ATG1a. (**C and D**) Transphosphorylation activity of ATG1a towards MBP (**C**) and MBP-ATG6 (**D**). *In vitro* kinase assay was performed with immunoprecipitated YFP-ATG1a in the presence of purified MBP and MBP-ATG6 proteins [shown in (**A**)]. The autoradiograph image shows that MBP was not phosphorylated (**C**), but MBP-ATG6 was phosphorylated (**D**) by YFP-ATG1a kinase. Coomassie blue (CBB) staining shows the input levels of MBP (**C**) and MBP-ATG6 (**D**), respectively. (**E**) Schematic representation of the kinase assay workflow for Figure 7 A-D. 4-leaf stage *35S:YFP-ATG1a* line seedlings were used for virus-induced gene silencing of glycolysis enzymes. After 10 ∼ 14 days, proteins were extracted from harvested leaves. YFP-ATG1a proteins were immunoprecipitated by GFP-Trap beads from each gene silenced sample. After brief washing, YFP-ATG1a and recombinant MBP-ATG6 (or MBP) proteins were used for kinase assay.

## Supplemental methods

### Yeast two Hybrid asasay

The entry clones containing *HXK1, CyTPI, GAPC1* or *PGK3* were subcloned into the destination vector pDEST-GADT7 (GAL4 activation domain) or pDEST-GBKT7 (GAL4 DNA binding domain). The *GAPC2* and *ATG101* entry clones were subcloned into the pDEST-GADT7 and pDEST-GBKT7 vectors, respectively. Yeast two hybrid assay was performed using Y2H Gold yeast strain, according to the manufacturer’s manual (Clontech). Protein-protein interactions were verified by growth in 3-out media (-leucine/-tryptophan/-histidine) containing 0.1 mM 3-amino-1,2,4-triazole (3-AT).

### Co-immunoprecipitation

The entry clones containing *HXK1, CyTPI, GAPC1, GAPC2* and *PGK3* were subcloned into the destination vector pEarleygate 201 for HA tagging, and the entry clone containig *ATG101* was subcloned into the destination vector pEarleygate 203 for c-myc tagging. *Agrobacterium C58C1* strains containing each destination vector and 35S:p19 construct were coinfiltrated into four-week-old *N. benthamiana* leaves at the OD_600_ adjusted to 1.0:1.0:1.5 ratio. At 48 h after infiltration, leaves were ground and mixed with an equal volume of ice-cold IP buffer (50 mM Tris-Cl pH 7.5, 150 mM NaCl, 10% glycerol, 5 mM EDTA, 1 mM DTT, 1% Triton X-100, 2 mM Na_3_VO_4_, 2 mM NaF, 20 mM β-glycerophosphate, and cOmplete protease inhibitor cocktail [Roche, 11836170001]). After centrifugation to eliminate cell debris, the supernatant containing 1 mg of total protein was incubated with EZview Red anti-c-myc affinity gel (10 μL gel per 1 mg of total proteins, Sigma-Aldrich, E6654) at 4℃ for 4 h. After incubation, the gel was washed four times with IP wash buffer (50 mM Tris-Cl pH 7.5, 150 mM NaCl, 5 mM EDTA, and 0.1 % Triton X-100). After washing, the gel was resuspended into 2× Laemmli Sample buffer. Ten μg of INPUT protein (1%) and 15 μL of IP elute were subjected to 10% SDS-PAGE. Immunoblotting was performed using anti-c-myc-peoxidase (Roche, 11814150001; 1:10,000) and anti-HA-peroxidase (Roche, 12013819001; 1:10,000). Signals were deteced by Imagequant LAS 4000 (GE Healthcare Life Sciences).

